# Identification and characterization of the antigonococcal prophage-encoded endolysin Phi1gp518

**DOI:** 10.64898/2026.07.09.737475

**Authors:** M. Pełka, B. Maciejewska, Z. Drulis-Kawa, A. Kwiatek, M. Adamczyk-Popławska

## Abstract

Gonorrhea, caused by the Gram-negative bacterium *Neisseria gonorrhoeae*, poses a growing global public health threat due to the rapid emergence of multidrug-resistant strains and the limited availability of effective treatments. Since there are no known lytic gonophages, we explored prophages present in the genome of *N. gonorrhoeae* FA1090, with a particular focus on prophage-encoded endolysins.

In this study, we evaluate antigonococcal properties of prophage-encoded endopeptidases with the NlpC/P60 enzymatic domain. Recombinant endolysin Phi1gp518 exhibits intrinsic bactericidal activity against non-permeabilized *N. gonorrhoeae* FA1090 cells. Furthermore, it shows an expanded host range against clinical gonococcal isolates. The gonolysin remains stable across all human body temperatures, a pH range of 5–10, and shows no cytotoxic effects toward human cervical epithelial cells, supporting its potential safety for therapeutic applications. Additionally, Phi1gp518 impairs the formation of gonococcal microcolonies and prevents proper biofilm establishment.

The antigonococcal properties of Phi1gp518 endopeptidase make it a good candidate for further protein engineering and development as an alternative treatment strategy for drug-resistant *N. gonorrhoeae* infections.

## Introduction

Gonorrhea, the second most common STD (sexually transmitted disease) is a rising public health problem. *Neisseria gonorrhoeae*, a Gram-negative diplococcal bacterium, is the etiological agent of gonorrhea. It belongs to one of the few bacterial genera characterized by a constant state of natural competence, resulting in high variability among strains. This variability is the obstacle in developing an effective vaccine against gonorrhea (1). Most of the gonococcal infections are asymptomatic, although they are more difficult to diagnose accurately among women. An untreated infection in female patients can lead to pelvic inflammatory disease, ectopic pregnancy, or infertility, resulting from tissue damage caused by the innate immune response involving neutrophils. Among men, potential outcomes include disseminated infection and increased risk of infectious arthritis and endocarditis (2). Current treatment relies on ceftriaxone (third-generation cephalosporin) and azithromycin (second-generation macrolide). However, due to the recent increase in the emergence of multidrug-resistant gonococci, WHO has set an alert regarding the issue, indicating the need to research alternative treatments (3). One of the promising novel treatments for drug-resistant strains is the use of bacteriophages, as they act as natural enemies of bacteria.

Bacteriophages were discovered over a century ago, however the interest in their therapeutic antimicrobial potential was lost due to the dawn of the antibiotic era. Currently, phage therapy is drawing interest again as a promising alternative to conventional bacterial infection treatment, including gonococci (4). Beyond whole phage particles, phage-encoded antimicrobial proteins are increasingly recognized as attractive candidates for targeted elimination of bacterial pathogens. Antimicrobial phage-encoded proteins belong to different types: 1) depolymerases (highly effective in degrading bacterial capsule, making them most promising agents for biofilm removal and resensitizing bacteria to immune system response), and 2) lysins (peptidoglycan hydrolases) (5). The last class can be divided into endolysins (encoded within lysis cassettes) and VALs (virion-associated lysins). Endolysins degrade cell wall structure in various ways, depending on their catalytic activity: glycosidases (e.g. lysozymes), amidases or endopeptidases. They act in the last step of the phage replication cycle, as they disrupt the cell wall, leading to cell lysis and release of new virions (6). Such diverse potential against extracellular bacterial structures can serve as an extension to conventional treatment solutions.

Administration of antibiotics do have advantages over phages – like controllable and precise dosage and product homogeneity. The same attitudes apply to phage-encoded proteins, however, enzybiotics (phage-encoded enzymes with antimicrobial activity) hold their own unique characteristics. Unlike antibiotics, phage proteins (especially endolysins) do not trigger rapid development of microbial resistance mechanisms (7, 8). By this, enzybiotics emerge as innovative biotherapeutic agents with substantial potential to complement or enhance existing antibiotic treatments.

The issue of the inability to find a lytic bacteriophage targeting *N. gonorrhoeae* (9) as a source of gonolysins shifted our interest to gonococcal prophage-encoded proteins. Prophage sequences among the *Neisseria* genus are highly abundant and very diverse, as 659 prophage sequences (1 - 90 kb) were identified within 85 gonococcal genomes (10). These regions act as evolution drivers, since half of the genomic rearrangement events among gonococci are caused by prophage elements (11). *N. gonorrhoeae* FA1090 carries five dsDNA and four ssDNA prophage sequences in its genome (12). NgoΦ1 and NgoΦ2 are intact prophages that, upon induction, produce phage particles with Sipho-like morphology (13). Incomplete dsDNA prophages include NgoΦ3, NgoΦ4 and NgoΦ5 (13). Such diverse proviral loads can be a rich source of genes with potential enzymatic properties.

Here we identified two endolysins derived from NgoΦ1 and NgoΦ3 prophage sequences present in the *N. gonorrhoeae* FA1090 genome. These enzymes are later regarded as ‘gonolysins’, a term referring to their origin and nature. Both proteins are endopeptidases carrying the NlpC/P60 catalytic domain, which is widespread among peptidoglycan remodeling enzymes of Gram-positive and Gram-negative bacteria. These peptidases are involved in numerous cellular processes, including growth, cell division, defense, virulence, sporulation, and biofilm formation (14). In this study, we are investigating the external use of prophage-encoded Gram-negative NlpC/P60 peptidases as potential antimicrobial agents against gonococci.

## Results

### Bioinformatic analysis of prophage-encoded endolysins

Identification of prophages within the *N. gonorrhoeae* FA1090 genome (GenBank: AE004969.1) was done originally by Piekarowicz et al., 2007 (13) and confirmed with modern bioinformatic tools by Orazi et al., 2022 (10). We reannotated the identified prophage sequences of the *N. gonorrhoeae* FA1090 genome with PHASTEST (15) and BV-BRC (16) online software. For further analysis, we took into consideration only potentially enzymatic proteins. Function prediction of proteins encoded by selected phage genes was determined with BLASTp (17), HHPred (18, 19) and Motif Finder (Kyoto University Bioinformatics Center) programs. After this step, two proteins were selected for further research – Phi1gp518 and Phi3gp1649. They are encoded within prophages NgoΦ1 and NgoΦ3, respectively. They are small (Phi1gp518 = 19.3 kDa, Phi3gp1649 = 19.6 kDa), highly similar proteins (88% of sequence identity), which vary at the first 51 aa of the N-terminal end. Main differences include an alteration in conformation and the presence of a hydropathic region (Fig. 1.). Both proteins carry a single enzymatic domain NlpC/P60, with peptidase activity, which cleaves the bond between D-Glu and mDAP (meso-diaminopimelic acid) present within the stem peptides of gonococcal peptidoglycan (14, 20). Domain identification confidence by BLASTp (e-value) is 1.08e-47 for Phi1gp518, and 8.67e-56 for Phi3gp1649. Structural comparison resulted in a TM-score of 0.965 (21) and RMSD of 1.24 Å (PyMol), indicating also very high (< 2Å) protein conformation similarity. Predicted structures of both potentially enzymatic proteins were solved with AlphaFold Server (22) and visualised with PyMol (The PyMOL Molecular Graphics System, Version 3.0 Schrödinger, LLC.).

**Fig. 1.**
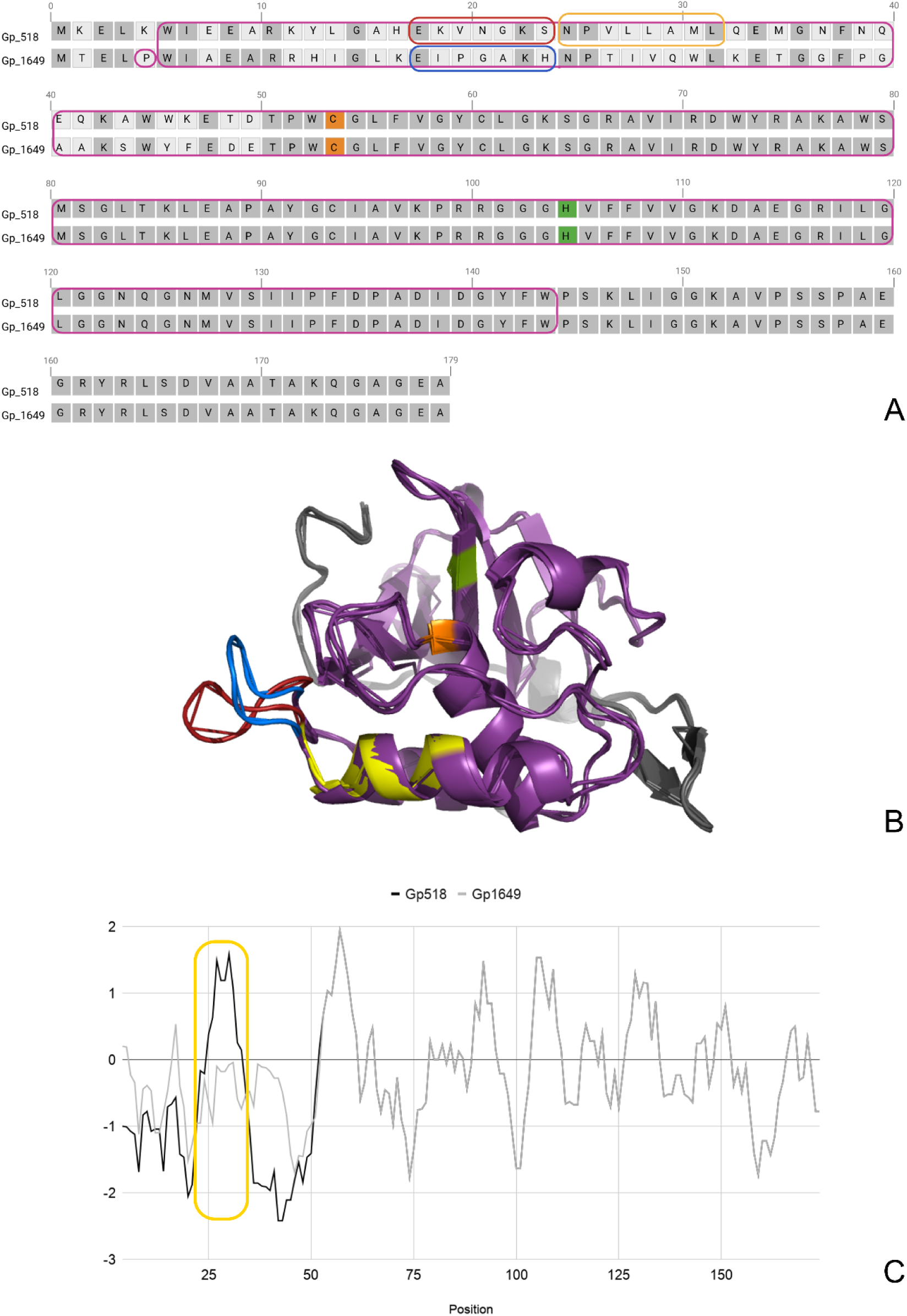
Visual comparison between Phi1gp518 and Phi3gp1649: **A** Sequence alignment with frames marking NlpC/P60 domain (purple), stronger hydrophobic site in Phi1gp518 (yellow), conformation difference between Phi1gp518 and Phi3gp1649 (red and blue), and predicted active site (orange and green); **B** 3D model of aligned proteins, with features marked the same colors as in figure 1A; **C** Hydropathy plot (Kyte & Doolittle scale) against amino acid positions with key difference between proteins marked with yellow box.

Cysteine peptidases of the NlpC/P60 family typically carry a catalytic dyad with a nucleophile and base residue (14). Active site prediction was done by identifying the canonical catalytic center (23) in the structure of Phi1gp518 and Phi3gp1649. Within the enzymatic domain, we found an N-terminal cysteine on the second α-helix and a histidine on the second β-strand. Selected possibly catalytic residues aligned with identical amino acids of Phi1gp518 and Phi3gp1649 sequence, indicating that Cys54 + His105 form a potential active site of the described endolysins (Fig. 1.).

### Identification of proviral lysis cassettes

To confirm that the selected proteins are endolysins (based on their genetic context), we identified potential lysis cassettes in the intact NgoΦ1 prophage and truncated NgoΦ3 prophage. Identical genes *ngo_0519* and *ngo_1650* are localized seamlessly downstream of *ngo_0518* and *ngo_1649*, respectively. Proteins encoded by genes *ngo_0519* and *ngo_1650* were analyzed with the TMHMM 2.0 online tool, which revealed that they both carry an N-terminal single transmembrane domain. Their relatively small size (12.7 kDa), structure, and genetic context suggest that proteins Phi1gp519 and Phi3gp1650 are holins. In addition, a set of two overlapping, small proteins (4.8 and 6 kDa) is encoded by identical pairs of genes *ngo_0520* + *ngo_0521* and *ngo_1651* + *ngo_1652*. These genes are localized downstream of endolysin and holin. They potentially encode o-spanin (identified by BLASTp with an e-value of 2e-24) and i-spanin (with a transmembrane central domain). These findings suggest that Phi1gp518 and Phi3gp1649 are part of lysis cassettes consisting of an endolysin, holin, and spanins (Fig. 2.).

**Fig. 2.**
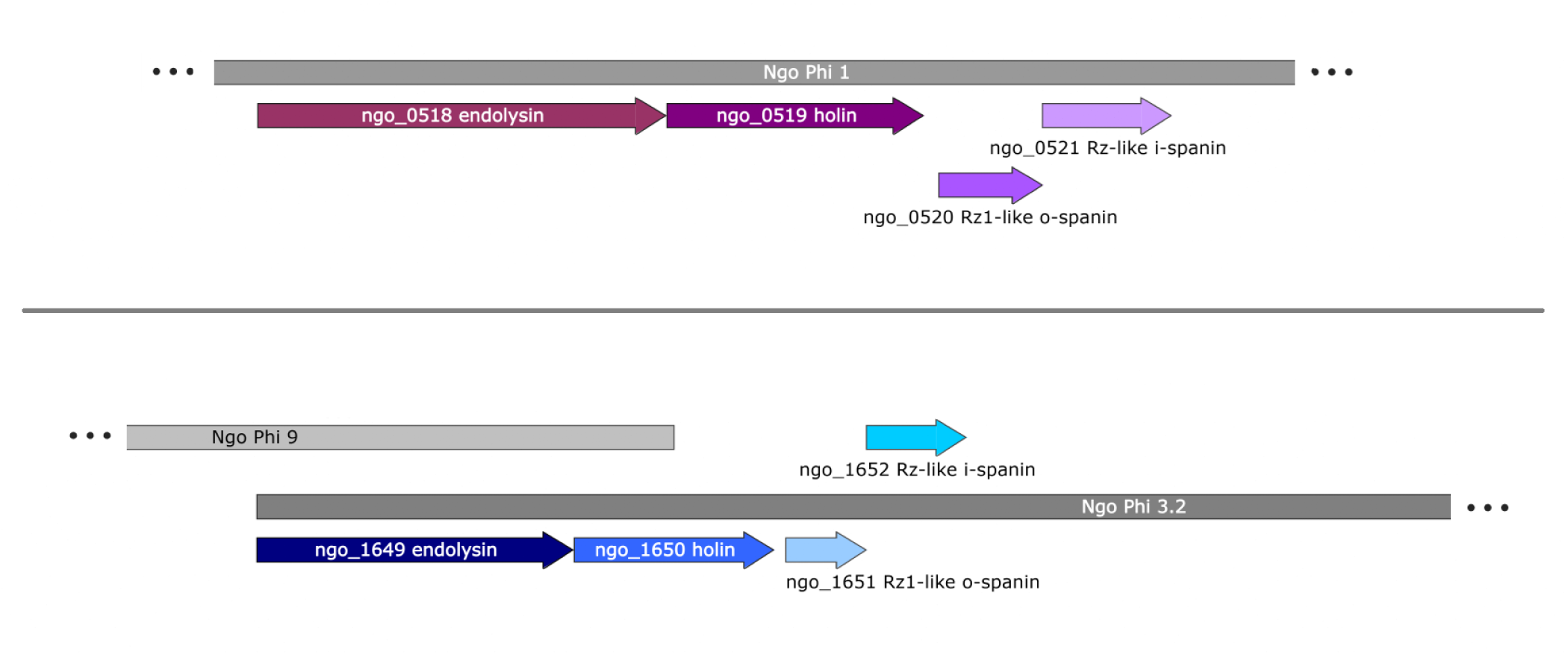
Putative lysis cassettes of prophages NgoΦ1 and NgoΦ3 from *N. gonorrhoeae* FA1090; NgoΦ3.2 is the second part of NgoΦ3, disrupted by NgoΦ9 ssDNA prophage; genes encoding endolysins and holins are seamlessly connected, and genes encoding spanins are overlapping.

### NlpC/P60 peptidases among *Neisseria* genus

The NlpC/P60 protein family is widespread among peptidoglycan hydrolases in both Gram-positive and Gram-negative bacteria (14). This family appears mostly in cell wall remodeling enzymes, which are involved in processes like cell growth and division. The maintenance of two highly similar NlpC/P60 peptidases within the *N. gonorrhoeae* FA1090 genome prompted our interest to investigate occurrence of them among *Neisseria* genus. We found cysteine peptidases similar to Phi1gp518 (Acc. No.: WP_010951071.1) and Phi3gp1649 (Acc. No.: WP_010951317.1) in other *Neisseria* genomes available in the GenBank database, including other *N.gonorrhoeae* strains, *N. meningitidis* and six commensal strains (Table 1). Most of the species possessed multiple NlpC/P60 peptidases. In order to evaluate the level of similarity between identified proteins, we performed sequence similarity analysis using BLASTp, multiple alignment with MAFFT (21), and conducted a structural comparison (Table 1., Fig. S1.). All proteins similar to Phi1gp518 (with TM-score > 0.95) were viral hits, which suggests that NlpC/P60 protein homologs from screened *Neisseria* genomes could be acquired through transduction by bacteriophages. Interestingly, BLAST database viral hits for NlpC/P60 proteins, except for Phi1gp518 and the one from *N. bergeri*, also show high similarity (100% cover, 70 – 93% identity) to the same phage-associated protein (Acc. No.: DAR31430.1) found in the human viral metagenome. Our bioinformatic analysis showed that this protein also displays the NlpC/P60 domain, but there is no experimental data, as this protein was identified in a computational metagenomic study (25). Overall, these findings underscore the importance of proviral load among *Neisseria* genus and confirms abundance of NlpC/P60 peptidases.

**Tab. 1.**
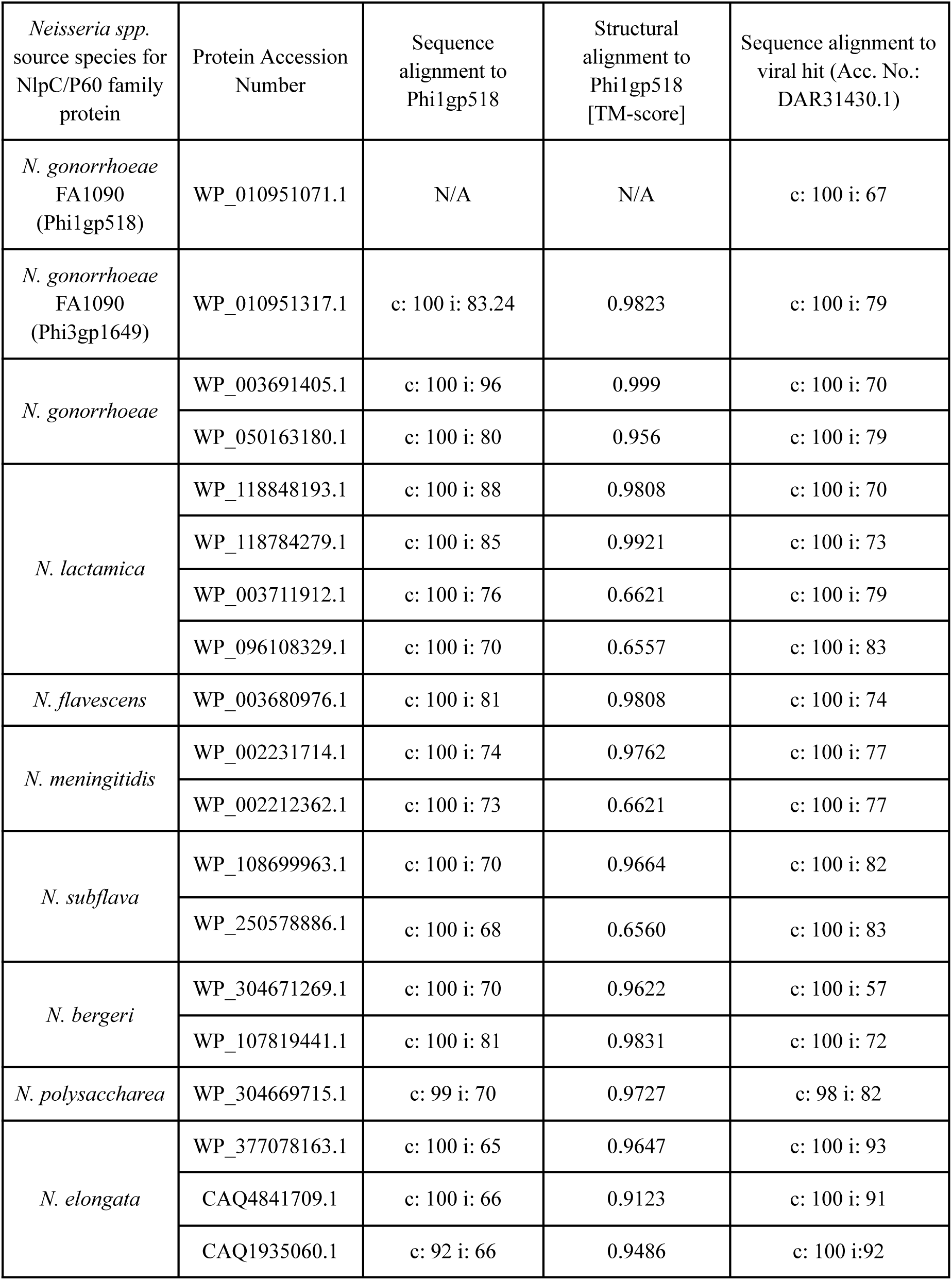
Identification of similarity level between NlpC/P60 family proteins among *Neisseria* genus; c: % cover, i: % identity.

### Purified prophage-encoded endolysins show muralytic activity

Potentially enzymatic proteins Phi1gp518 and Phi3gp1649 were overexpressed with an additional 6x His-tag on the N-terminal end. Efficiency of gonolysin synthesis was high, as protein yield reached 32.6 or 26.6 mg per liter of liquid culture for purified Phi1gp518 and Phi3gp1649, respectively.

For biochemical confirmation of gonolysin muralytic activity, we performed a zymogram assay with lyophilized *M. lysodeikticus* cells as a substrate. To prevent false positive results, one zymogram gel was stained without the renaturation step (24). This allows distinguishing gel destaining caused by biochemical properties from the actual lytic ability of the protein. For this reason, the lysozyme effect was considered a false positive, as it produced clear bands on both gels. In contrast, mutanolysin, used as a positive control, yielded clearance only on the renatured gel, and the same pattern was observed for the two tested proteins. Clear bands on the renatured zymogram demonstrated that both Phi1gp518 and Phi3gp1649 exhibit muralytic activity at 1 mg/mL against *M. lysodeikticus* peptidoglycan (Fig. 3.1, lanes A, B).

**Fig. 3.**
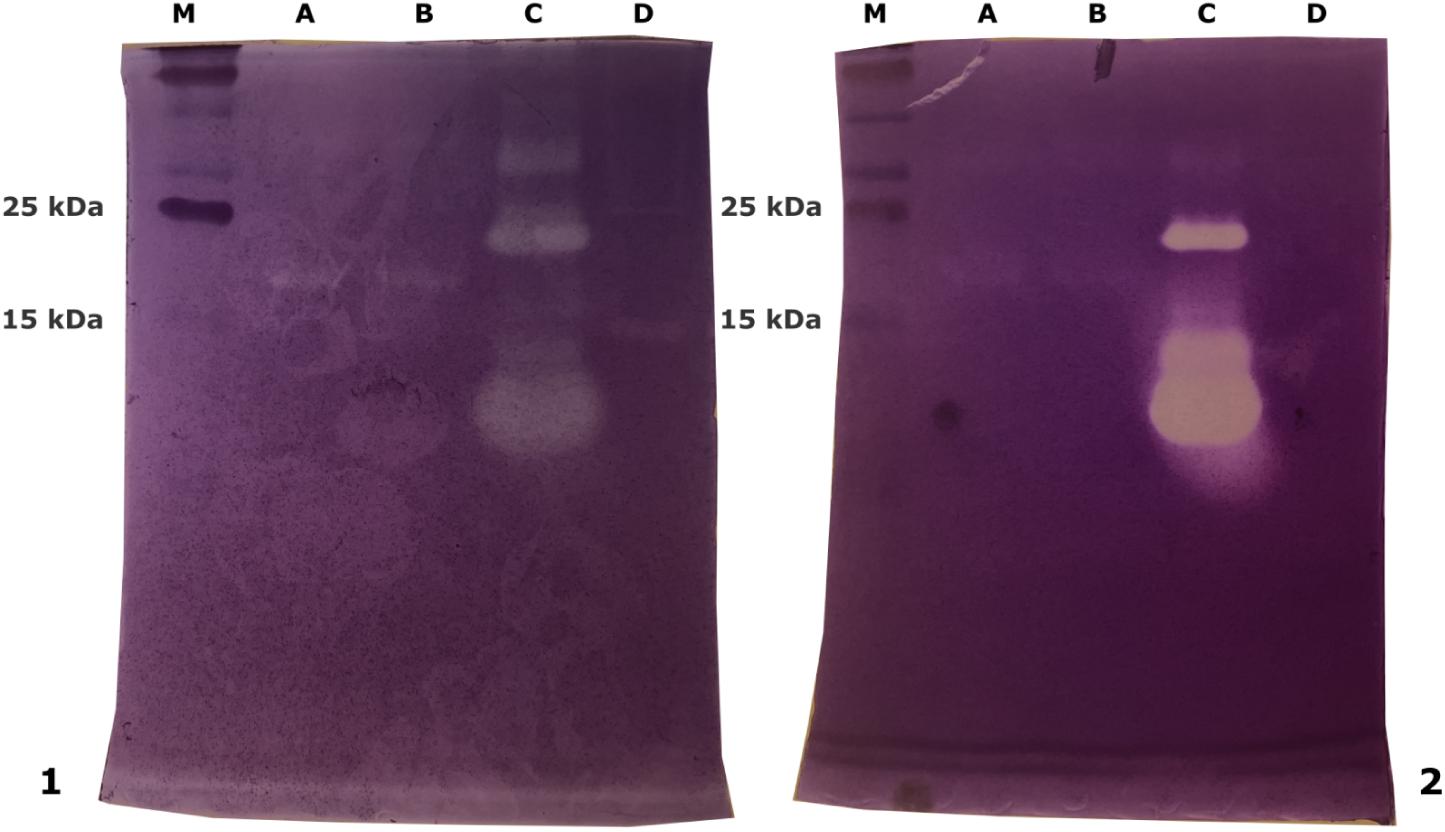
Zymogram assay with 10% of *M. lysodeikticus* peptidoglycan in an SDS-PAGE gel; samples on the (1) renaturated or (2) non-renaturated gel are: marker (M), Phi1gp518 (A), Phi3gp1649 (B), lysozyme (C), mutanolysin (D)

### Antimicrobial activity of endolysins on a pathogenic gonococcal strain

In general, endolysins do not exert bactericidal activity against Gram-negative bacteria because their access to the peptidoglycan layer is hindered by the outer membrane. First, we tested the antigonococcal effect of Phi1gp518 and Phi3gp1649 on permeabilized gonococci (after incubation in chloroform-saturated PBS to punctuate the outer membrane) using the opacity test in a glass tube for 30 min. Initial *in vitro* effect of both purified endolysins could be observed within the first 2 min of incubation on the bench (Fig. 4.).

**Fig. 4.**
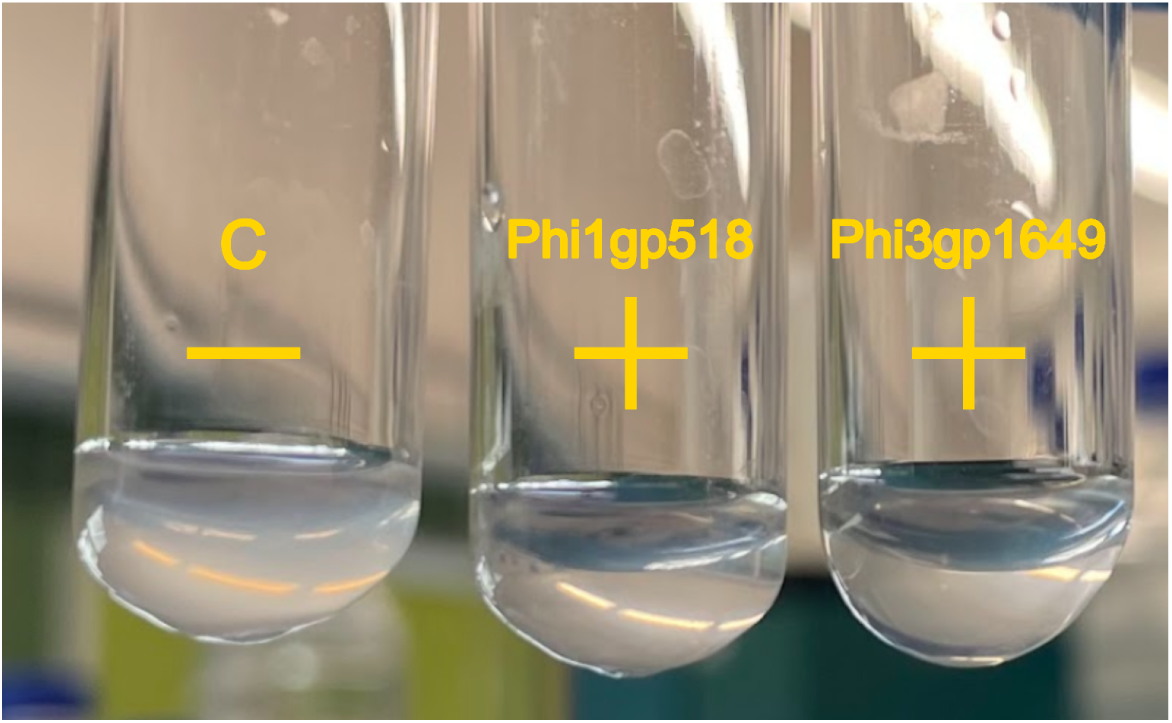
Opacity test of Phi1gp518 and Phi3gp1649 against permeabilized gonococci; C – untreated bacteria; ‘–’ and ‘+’ indicate negative or positive test result, respectively.

Second, we adapted the turbidity reduction assay (TRA) (27) to *N. gonorrhoeae* FA1090. The preliminary TRA test measured the enzymatic activity of Phi1gp518 and Phi3gp1649 added to permeabilized bacteria at the final concentration of 500 nM or 1000 nM. Although both endolysins reduced the optical density of bacterial culture over time, only Phi1gp518 showed hallmark enzyme-like kinetics with burst phase and plateau saturation point (Fig. 5.). Since our research interest at the time was to identify an effective anti-gonococcal enzyme, further research was done on Phi1gp518.

**Fig. 5.**
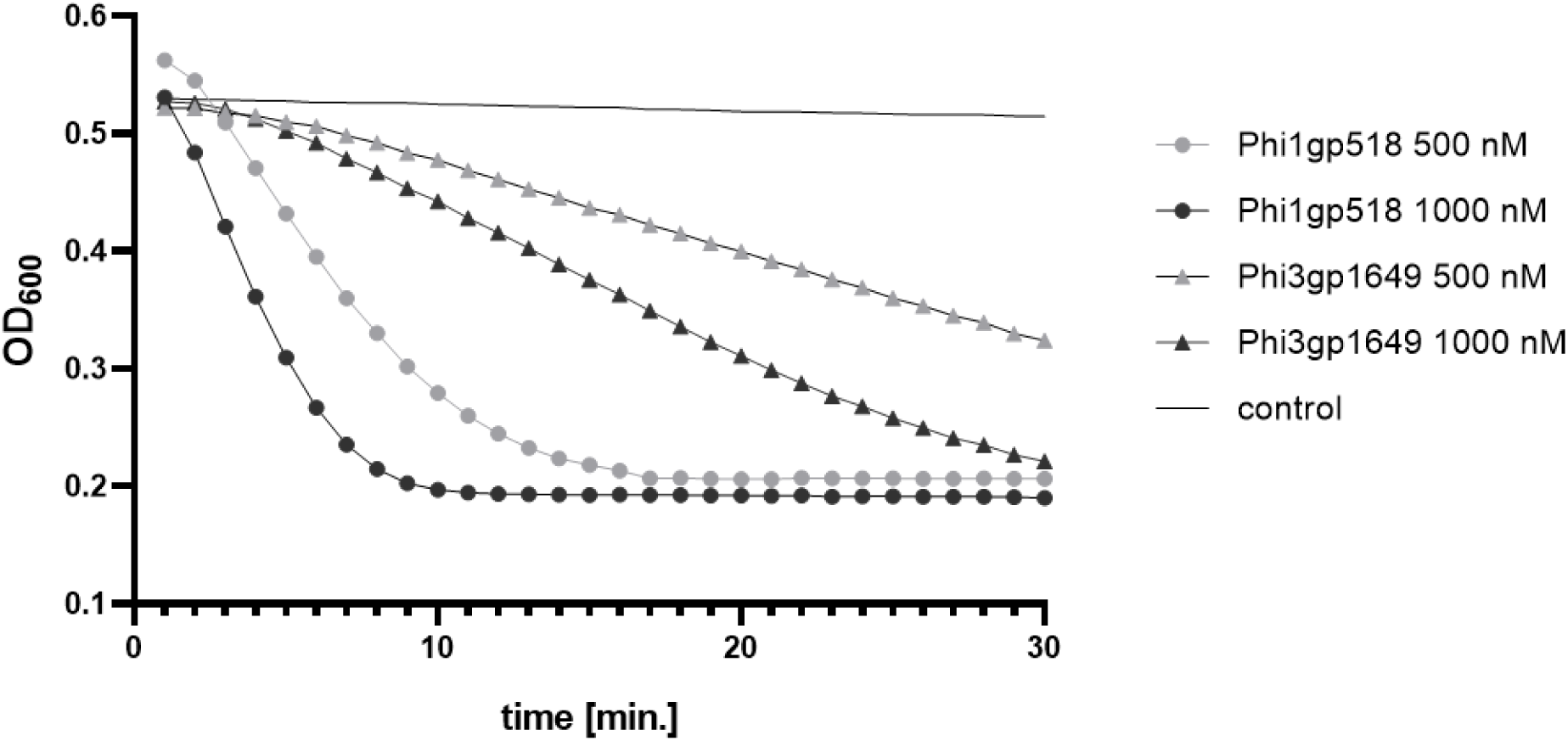
TRA test results for the lysis effect of Phi1gp518 and Phi3gp1649 purified proteins on permeabilized *N. gonorrhoeae* FA1090 cells

Using the standardized endolysin activity calculation method developed by Briers et al., 2007 (28), we established a slope ΔOD_600_ / min curve plotted against a range of Phi1gp518 concentrations (Fig. 6.). The calculated enzyme activity was 548 U/mg (R^2^ = 0.99).

**Fig. 6.**
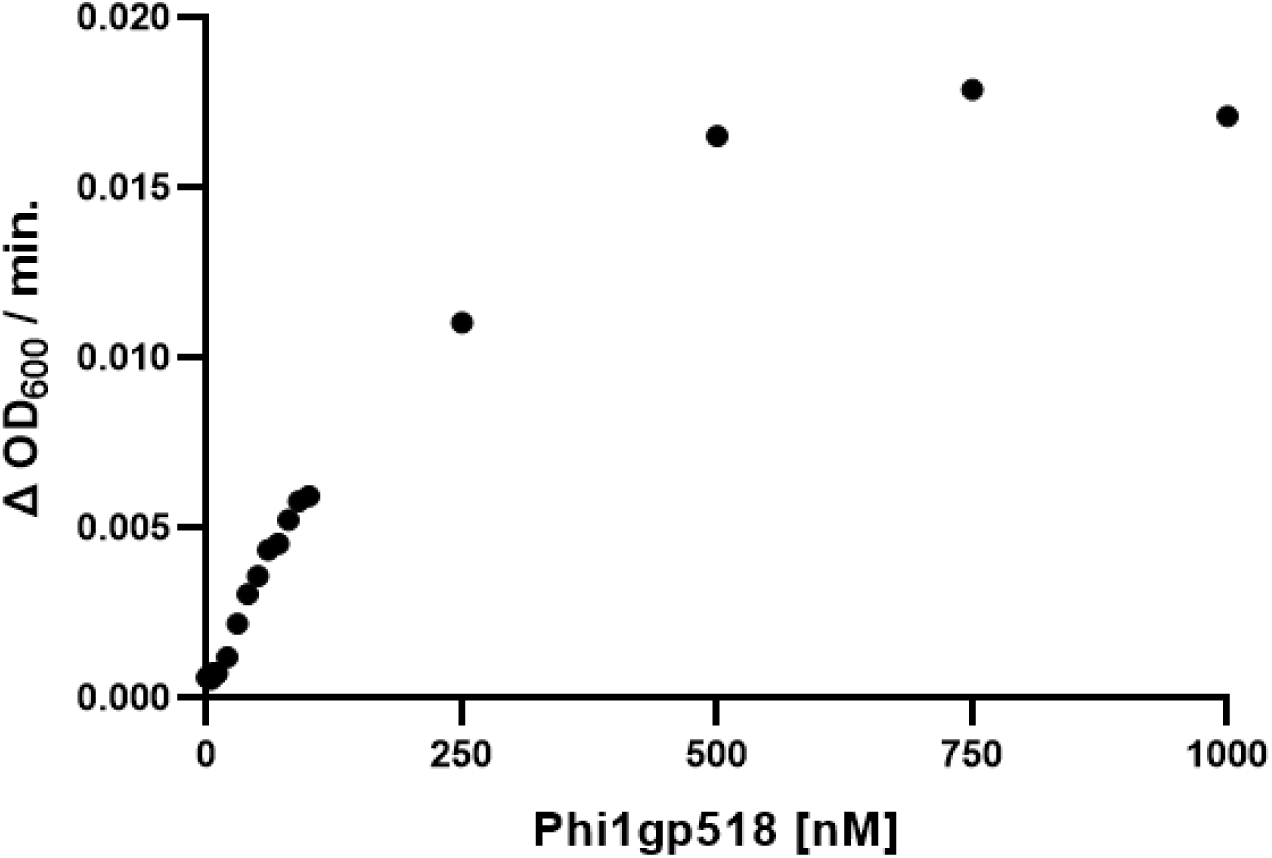
Phi1gp518 enzymatic activity against *N. gonorrhoeae* FA1090 at 1 – 1000 nM concentration of purified enzyme; curve equation for linear region (first 11 points) is *2.74x + 3.5e-4*

### Effect of pH and temperature on gonolysin Phi1gp518 activity against permeabilized *N. gonorrhoeae* FA1090

To further characterize Phi1gp518, we evaluated by TRA assays how exposure to different pH levels and temperatures affects its efficiency. The enzyme remained stable (> 85% relative activity) in PBS or Tris-Cl buffers with a pH level from 7 to 10. The highest activity was observed at pH 8 (Fig. 7A). Relative enzymatic activity >85% was maintained at room temperature, as well as in the range 30–42°C, with 37°C being the optimum (Fig. 7B).

**Fig. 7.**
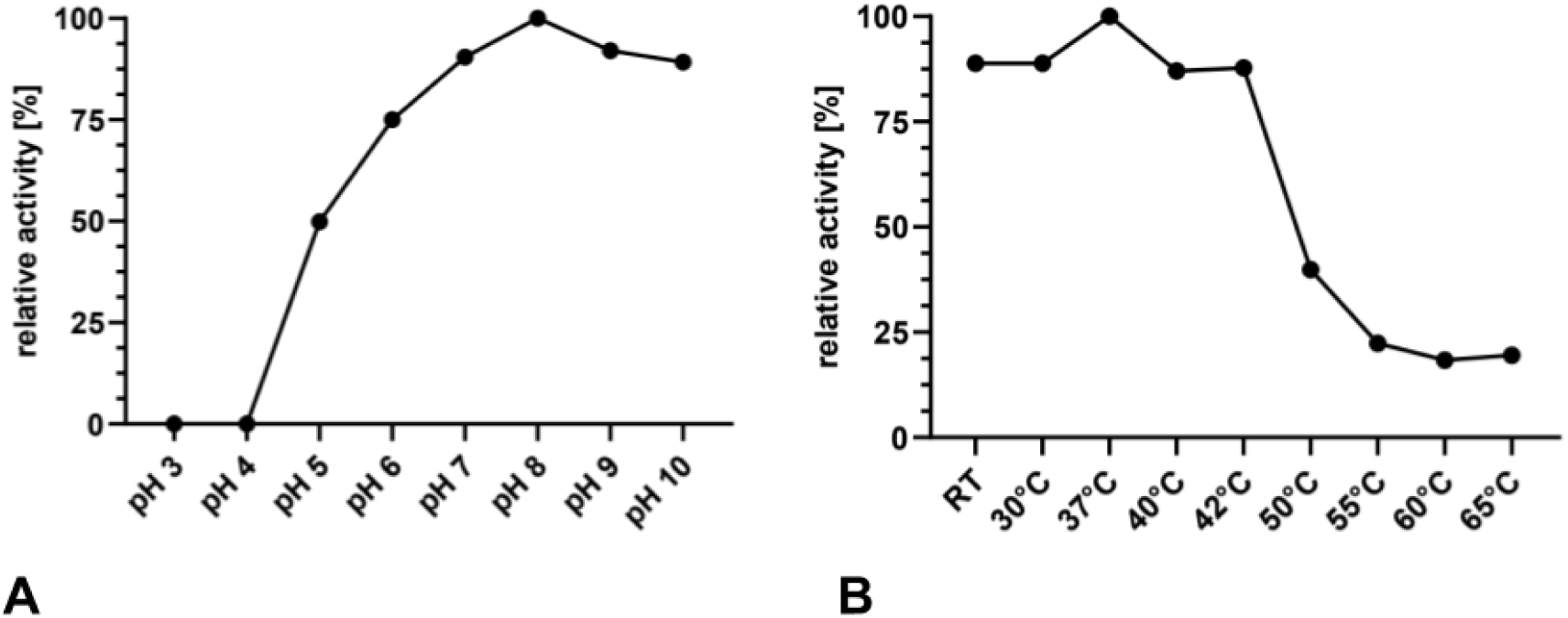
Activity of Phi1gp518 on permeabilized *N. gonorrhoeae* FA1090 (A) incubated at a range of pH levels and (B) after preincubation at different temperatures; relative activity was calculated as the total decrease of OD_600_ in 45 min relative to the value with highest performance.

### Phi1gp518 shows intrinsic enzymatic activity against non-permeabilized gonococci

The presence of the hydrophobic region in Phi1gp518 may suggest partial accessibility to intact cells and intrinsic enzyme activity (29). To assess it, we performed a comparative TRA test on permeabilized and non-permeabilized (later regarded as ‘raw’) *N. gonorrhoeae* FA1090 cells. Gonolysin treatment resulted in a 3-fold difference in absolute OD_600_ decrease (background-corrected) between permeabilized and raw bacteria, measured over 45 min. Although observing a decline in the ability to lyse raw bacteria, the comparison of treatment against control was still significant (Fig. 8.). Gonolysin treatment resulted in a ΔOD_600_ decrease of average 0.316 in permeabilized cells and 0.148 in raw cells, compared to 0.023 and 0.05 in the respective controls.

**Fig. 8.**
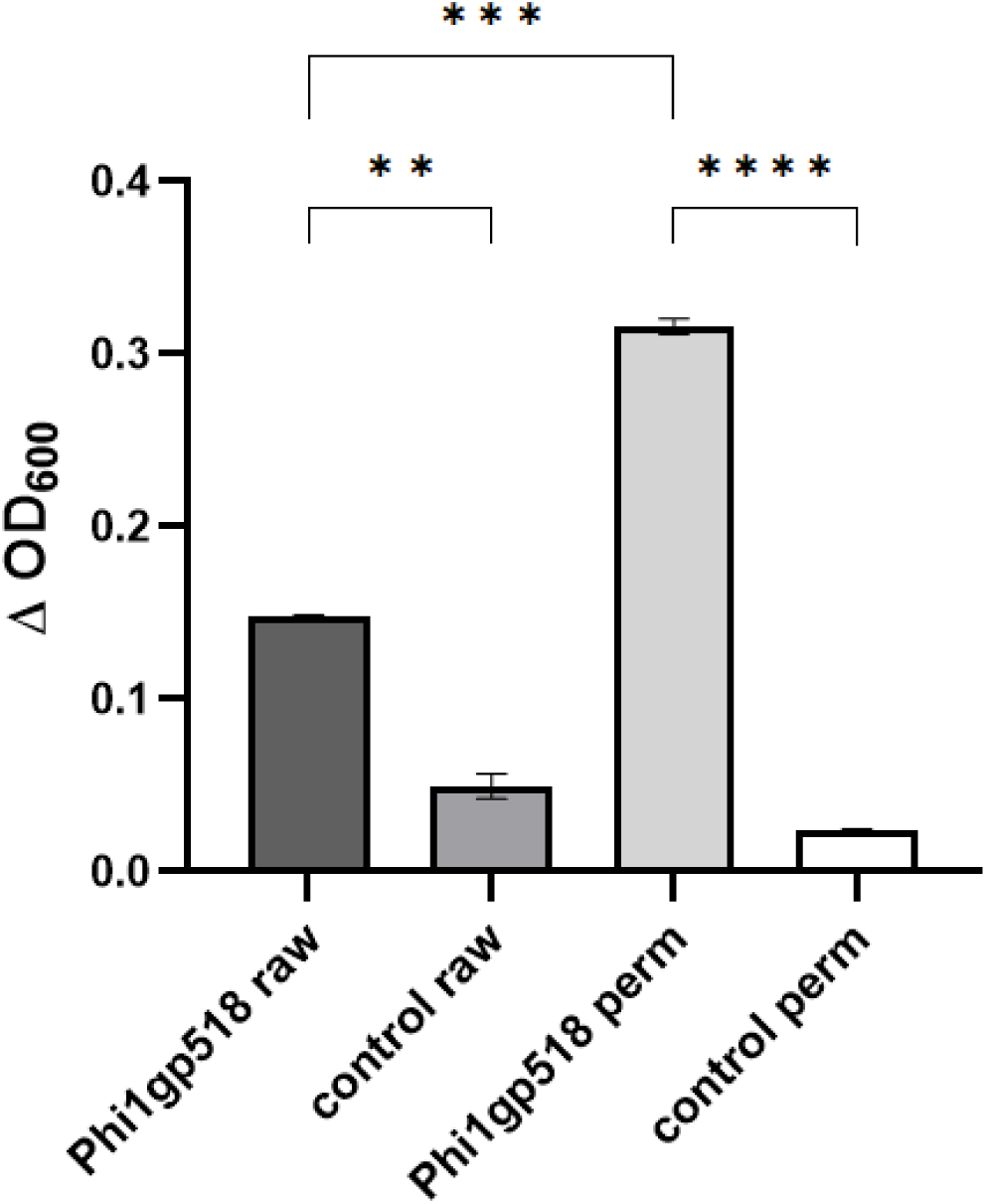
Phi1gp518 activity on permeabilized or raw *N. gonorrhoeae* FA1090 cells; statistical significance is marked with ** for p-value <0.01, *** for p < 0.001 and **** for p < 0.0001; Δ OD_600_ value was calculated as difference between OD_600_ at starting and ending time point of measurement.

### Diverse specificity of Phi1gp518

To assess the Phi1gp518 lytic spectrum, we performed a modified TRA assay on different, non-permeabilized bacterial species (Tab. 3.). In addition to clinical *N. gonorrhoeae*, we tested *E. coli*, *L. plantarum,* and *G. vaginalis* strains, as these species may be present (physiologically or pathologically) in the female urogenital tract (30). Gonolysin Phi1gp518 shows comparable OD_600_ reduction towards some of the raw gonococcal strains (FA1090, 077, β-) and both *E. coli* strains (Fig. 9A, Fig. S2.). The effect on other *N. gonorrhoeae* strains was more than 2-fold lower. Interestingly, the absolute OD_600_ value for *L. plantarum* and *G. vaginalis* increased over time. However, these results calculated as percentage of OD_600_ loss gives a more detailed understanding of enzyme activity on a given strain. From this point of view, Phi1gp518 seems to show highest specificity towards the *N. gonorrhoeae* FA1090 strain (Fig. 9B).

**Fig. 9.**
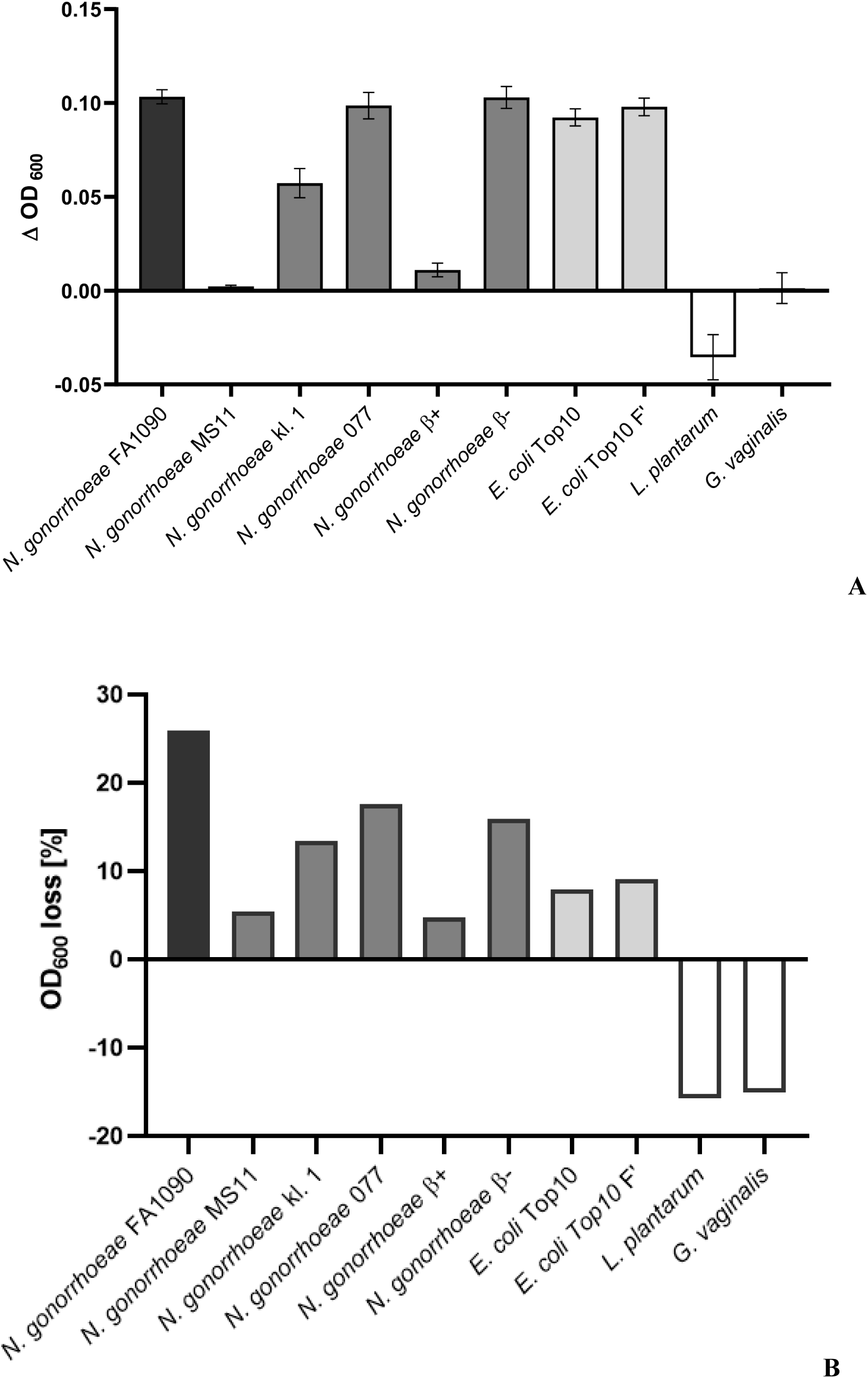
(A) Changes in OD_600_ determined by TRA test on different raw bacterial Gram-positive and -negative vaginal strains; Δ OD_600_ value was calculated as the difference between OD_600_ at the starting and ending time point of measurement, with subtraction of the control (untreated bacteria) value for each strain; (B) % of OD_600_ loss calculated as: *(OD start - OD stop)/ OD start*

### Phi1gp518 limits the formation and surface attachment of gonococcal biofilm

During preliminary experiments, a viable biofilm with permeabilized cells was hard to maintain, as this process weakens the cells. Moreover, our data showed that *N. gonorrhoeae* is sensitive to many commonly used OMPs (outer membrane permeabilizators). For these reasons, we conducted biofilm experiments without permeabilizing the cells, as Phi1gp518 showed intrinsic membrane penetration ability. The crystal violet staining assay first measured the ability to disrupt a gonococcal biofilm. Gonococcal biofilm biomass, measured as OD_600_ value, was significantly reduced for all tested Phi1gp518 concentrations (Fig. 10.). Application of 3000 nM Phi1gp518 resulted in a 1.5-fold decrease compared to 1000 nM. However, there was no substantial improvement between treatment with 3000 nM or 6000 nM of Phi1gp518.

**Fig. 10.**
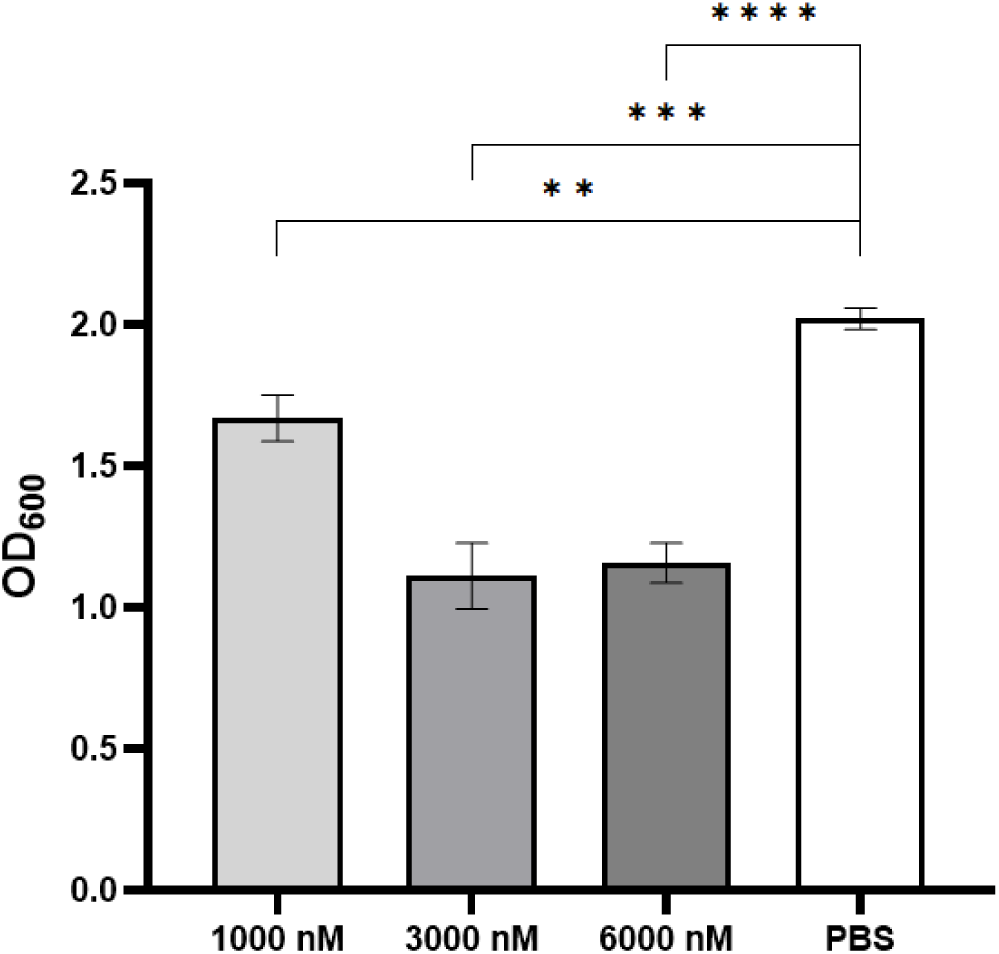
Gonococcal biofilm biomass measured with crystal violet staining; statistical significance is represented by asterisks: ** (p < 0.01), *** (p <0.001), or **** (p < 0.0001)

To better understand the influence of gonolysin on biofilm formation, we set up a plate with *N. gonorrhoeae* FA1090 cells, treated immediately with 3000 nM Phi1gp518 and incubated it statically for 12 h. After incubation time, in the control sample, bacterial cells were organized in visible clusters, forming microcolonies which are the basis for proper biofilm establishment. The gonolysin treatment successfully inhibited gonococcal microcolony formation by dispersing the cells (Fig. 11.).

**Fig. 11.**
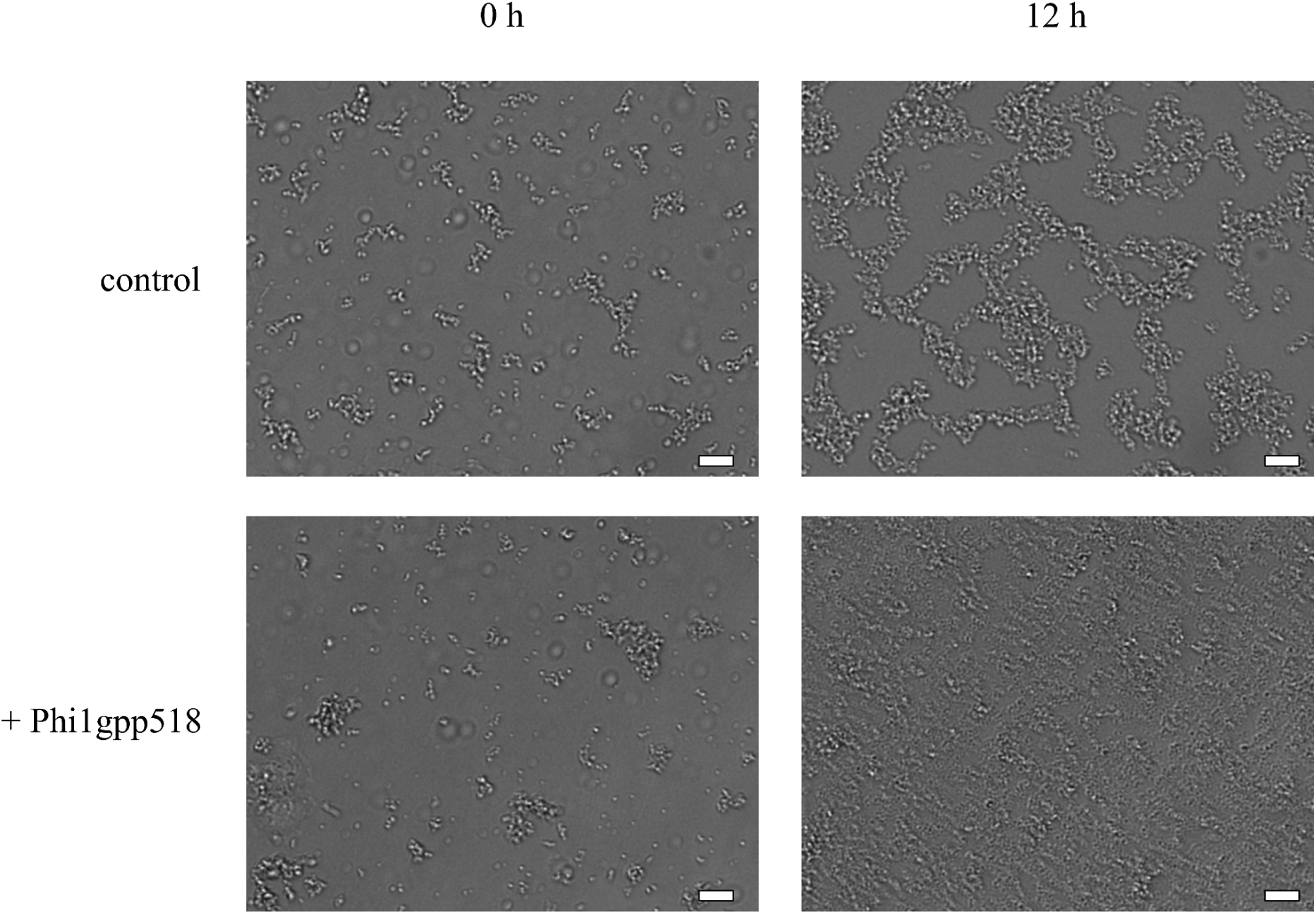
Microscopy bright light images from JuLi Stage ™ plate reader; *N. gonorrhoeae* FA1090 cells were treated with Phi1gp518. The scale bar is 20 μm.

To search for potential changes in gonococcal biofilm structure caused by endolysin treatment, we visualized biofilms by confocal microscopy using fluorescence staining. Treatment of the 20-hour biofilm with Phi1gp518 resulted in a twofold reduction in the live: dead ratio (0.47) compared to the control (1), as determined by integrated density factor using ImageJ software. Importantly, gonolysin treatment led to complete detachment of the biofilm biomass from the substratum, which is consistent with our previous observations, indicating that in the presence of gonolysin, bacteria were unable to form clusters on the substratum (Fig. 12.). The biofilm structure appeared more relaxed when formed in the presence of Phi1gp518.

**Fig. 12.**
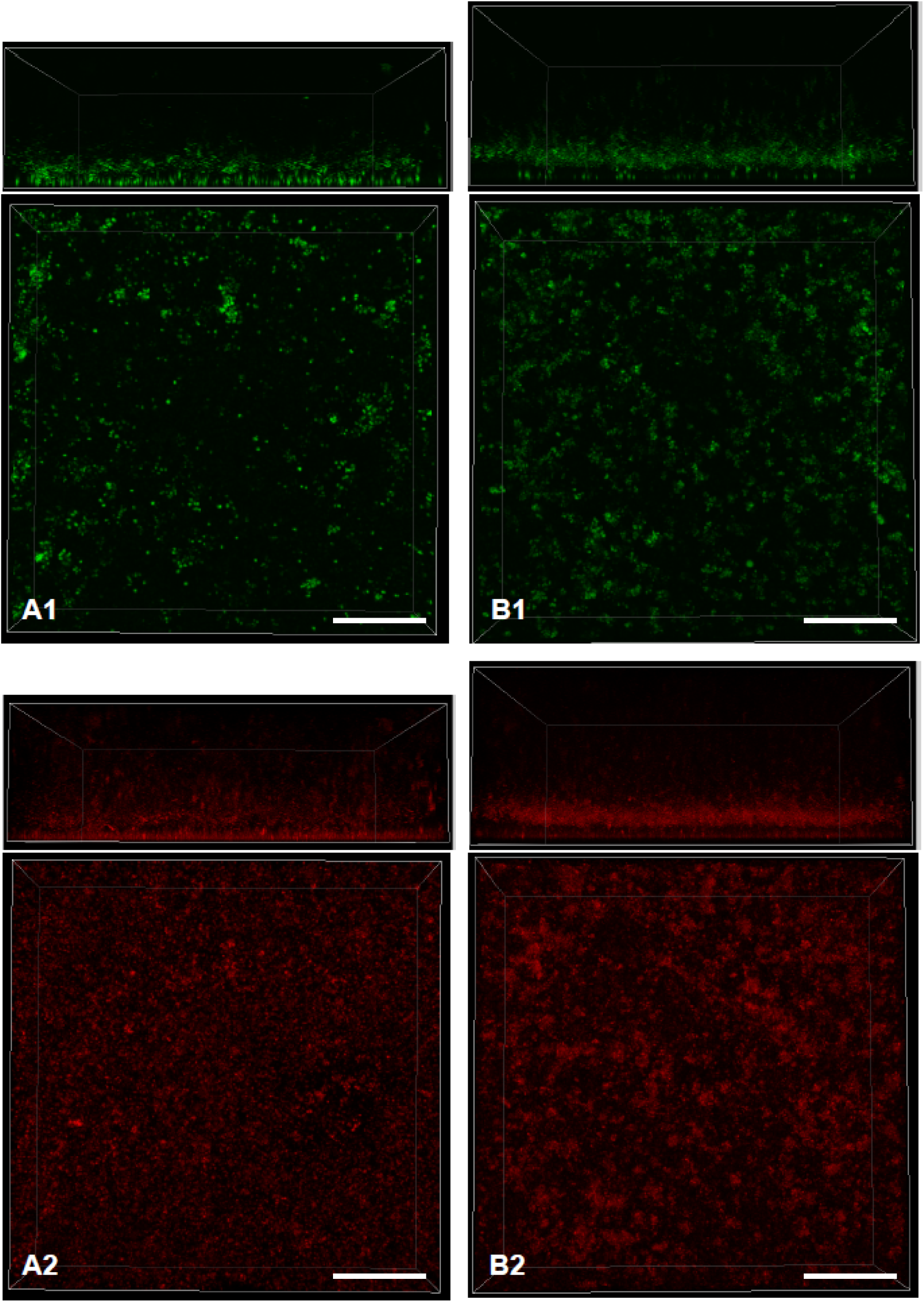
Confocal microscopy images of gonococcal biofilm treated with PBS (A) or 3000 nM Phi1gp518 (B) with visible live (1) and dead (2) cells. Bottom and side views; the scale bar is 200 μm

### Phi1gp518 gonolysin is not cytotoxic to cervical epithelial cells

The MTT cytotoxicity test was carried out on the ME-180 cell line to evaluate the impact of Phi1gp518 on the human cervical epithelial cells, most prone to be targeted during gonorrhea infection. Overall cell viability in treated samples remains >85% within the first 24 h (Table 2.). Cytotoxicity increases over time by 13.5% and 14.5% for 3000 nM and 6000 nM endolysin treatment, respectively. However, this effect can partially result from solvent composition, as the dialysis buffer contains a high (250 mM) NaCl concentration. Comparison to this control does not show statistical significance (p > 0.05), indicating that Phi1gp518 is not cytotoxic to ME-180 cells (Fig. 13.).

**Fig. 13.**
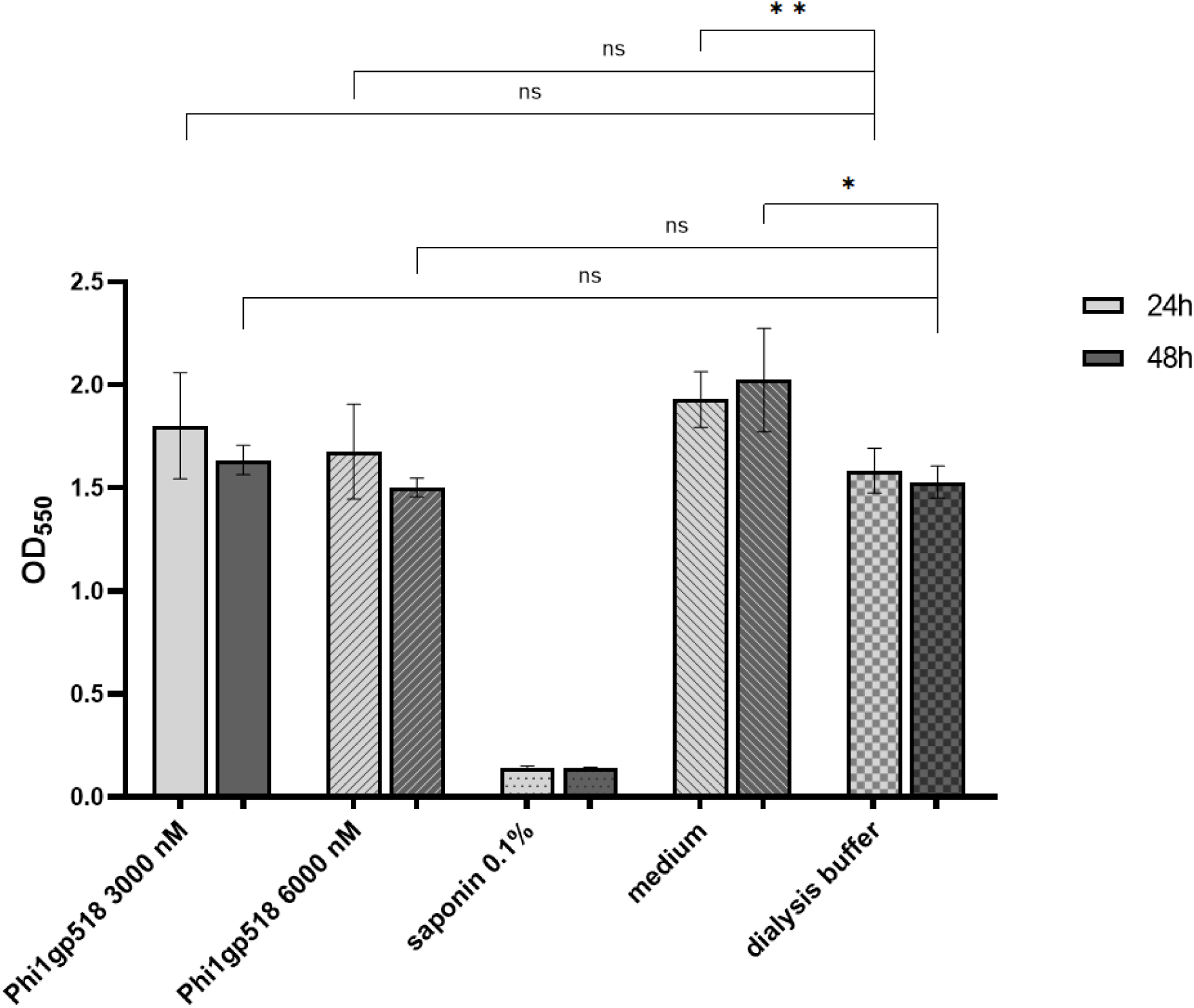
Mean OD_550_ values for MTT cytotoxicity evaluation on human epithelial ME-180 cell line with 0.1% saponin as positive control; statistical significance compared to the negative control (dialysis buffer) is represented by asterisks: * for p-value < 0.05, ** p < 0.01

**Tab. 2.**
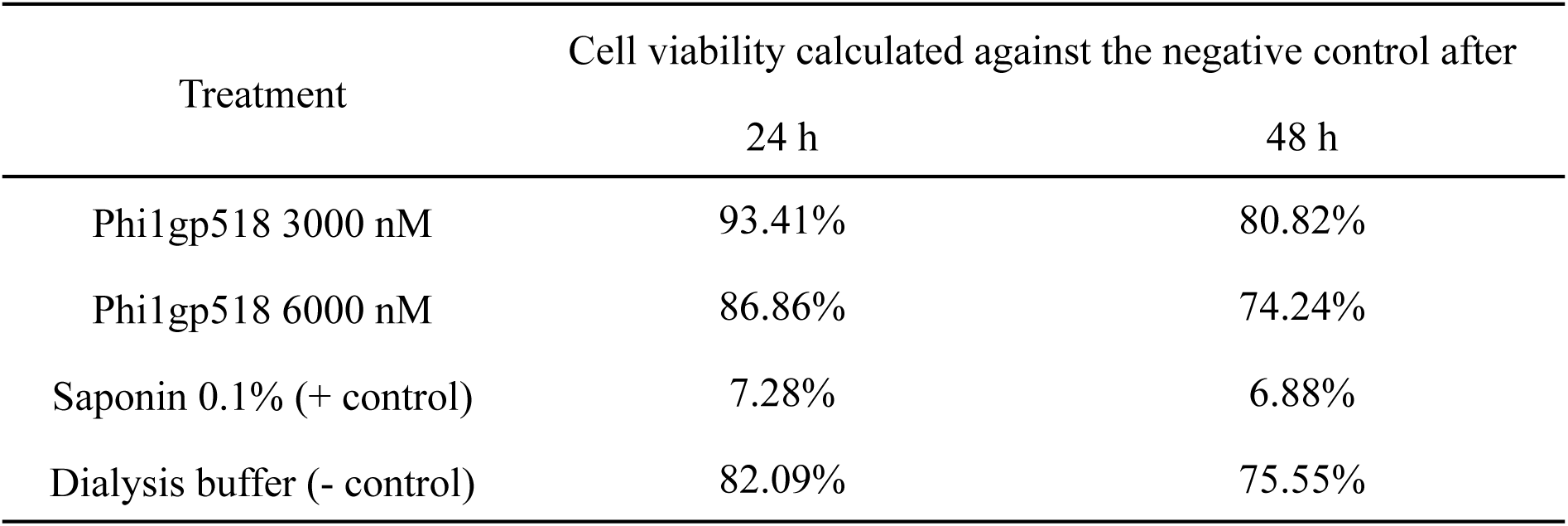
Cytotoxicity levels of Phi1gp518 towards human epithelial cells.

## Discussion

Gonorrhea remains a major global health concern due to increasing antimicrobial resistance of *N. gonorrhoeae* strains and the limited availability of alternative therapeutic strategies. In contrast to many other bacterial species, no lytic bacteriophages infecting *N. gonorrhoeae* have been described to date (4, 9). This lack of known lytic gonophages prompted our interest in proviral sequences present within gonococcal genomes. Optimized viral genomes, shaped by selection pressure (31), make even partial prophage sequences a rich reservoir of diverse genetic material. Our data indicate that the total prophage gene density (expressed as CDS per Mb) is three times higher than in the rest of the *N. gonorrhoeae* FA1090 genome, which underscores prophage sequences contributing to 6.87 % of the total genome size (32). This comparison highlights the exceptional value of gonococcal prophages as a source of genes potentially encoding proteins with antimicrobial activities, effectively making them ‘genetic hotspots’ within their host genomes.

Numerous prophages have been identified in *N. gonorrhoeae* strains, including the widely used model FA1090. We screened *N. gonorrhoeae* FA1090 genome for prophage genes encoding potential lysins and identified at least three proteins containing a putative NlpC/P60 peptidoglycan hydrolase domain: Phi1gp508, Phi1gp518, and Phi3gp1649. This domain is present in papain-like proteases with conserved Cys and His at the catalytic site. In general, enzymes from the NlpC/P60 family are exploited by bacteria for cell wall remodeling, cell separation, and interactions with host organisms (14). Our preliminary analysis indicates that, within the *Neisseria* genus, NlpC/P60-containing enzymes are predominantly encoded by prophages (Table 1.). This suggests that gonococci may have acquired or enhanced their peptidoglycan remodeling capacity through lysogenic conversion. Notably, the lysis cassette was not disrupted by insertion of NgoΦ9 into the NgoΦ3 region, further supporting the functional relevance of these enzymes.

To date, only one active phage-encoded NlpC/P60 endolysin has been reported. It was identified in mycophage D29 as a multi-domain enzyme; however, the authors demonstrated highly specific anti-biofilm activity of the enzymatic domain alone (33). Since gonococci are characterized by relatively high peptidoglycan cross-linkage (34), an endolysin with peptidase activity (such as Phi1gp518) can potentially disrupt cell wall architecture efficiently. The NlpC/P60-containing endopeptidases (Phi1gp518 and Phi3gp1649) are highly similar in structure, and we investigated their properties as candidates for antimicrobial drug development. After preliminary TRA results demonstrated its high efficiency against gonococci, Phi1gp518 became the main focus of our study. Bioinformatic analysis also showed an amphipathic helix in the Phi1gp518 sequence (absent in Phi3gp1649), indicating a promising starting point for future engineering of the protein (35). As this region is possibly causing membrane permeability, its modifications might improve the intrinsic activity of Phi1gp518 against gonococci.

We demonstrated that external application of purified Phi1gp518, at high concentrations, exhibits antimicrobial activity against *N. gonorrhoeae* FA1090 cells. The observed effect of gonolysin treatment on biofilm formation suggests that this protein, acting on peptidoglycan, disrupts interactions between bacterial cells (preventing formation of microcolonies) and influences adhesion between bacterial cells and the substratum (leading to biofilm detachment). This may be the underlying cause of impaired biofilm development and total biofilm biomass reduction. Phi1gp518 remains stable at temperatures representing physiological or pathological human body states (30–42 °C), supporting both topical and internal administration, including severe infections. No significant loss of activity was observed at extreme pH values (5 – 10). The activity of gonolysin is therefore unlikely to be affected by pH levels at different body sites that can be infected with *N. gonorrhoeae*: genital tract (pH 3–5), throat and ocular mucosa (pH 6–7), or rectum (pH 7–10) (36–38). The protein itself, even at high concentrations, did not show cytotoxic activity towards human cervical epithelial cells.

Endolysins generally show limited activity against Gram-negative bacteria due to the presence of an outer membrane (6). We demonstrated that Phi1gp518 has an effect against ‘raw’, i.e., non-permeabilized *N. gonorrhoeae* FA1090 and other clinical strains. The variable effect among our collection of gonococcal strains underscores morphological differences of *N. gonorrhoeae*. One key difference among isolates may be the gonococcal capsule-like structure composed of polyphosphates (39). In fact, Phi1gp518 has a strong bactericidal effect against permeabilized clinical *N. gonorrhoeae* strains other than FA1090 (Fig. S3.). Activity towards *E. coli* strains indicates that Phi1gp518 has a wide lytic spectrum, which is considerable among Gram-negative endolysins (40). The sequence of pentapeptide (which is probably the substrate for Phi1gp518) in the *E. coli* cell wall is the same as gonococcal (41).

Importantly, incubation of *L. plantarum* and *G. vaginalis* with gonolysin resulted in a significant increase in OD_600_ over time. This effect is likely due to lactate secretion rather than cell proliferation, as the doubling time of these bacteria is > 100 min (42, 43) and changes in OD were observed for 45 min. Regardless, it indicates that gonolysin treatment has no negative impact on vaginal microbiota.

Live/dead staining of *N. gonorrhoeae* biofilm treated with Phi1gp518 did not reveal a significant bactericidal effect; instead, we observed total detachment of the biofilm from the substratum. Cells also seemed to be more dispersed, suggesting disruption of diplococcal morphology. Gonolysin impaired an early step of biofilm development, namely the formation of microcolonies (2). Although cell dispersion is part of the colonization process, sudden removal of whole biofilm biomass may disrupt biofilm homeostasis and limit bacterial accumulation (44).

One of the virulence strategies of *N. gonorrhoeae* involves the release of toxic peptidoglycan (PG) fragments that are recognized by intracellular receptors NOD1 and NOD2. To trigger NOD1 activation, the agonist must contain the second and third amino acids of the stem peptide or be a free full peptide (which requires amidase activity). NOD2 receptors, in contrast, recognize macromolecular PG fragments (45). However, no enzyme with proven endopeptidase activity has yet been identified as directly involved in the synthesis of toxic PG fragments (20). Based on its cleavage specificity, an NlpC/P60 peptidase is predicted not to release the canonical NOD1 ligand, as this enzyme hydrolyzes the bond between amino acids triggering the NOD1 response. Although this issue requires further research, the enzymatic activity suggests little potential cytotoxic effect of treatment with gonolysin.

In conclusion, we described the first endolysin targeting *N. gonorrhoeae*. Endolysins are promising candidates for novel antibiotic alternatives. Development of resistance mechanisms against endolysins is unlikely, as administering intracellular enzymes externally significantly lowers the risk of bacterial resistance. Moreover, variants resistant to endolysins often display reduced growth rates (due to mutations in vital structures such as peptidoglycan), and such mutations are frequently reversed (46). The intrinsic activity of Phi1gp518, likely resulting from its N-terminal hydrophobic region, makes this enzyme a good candidate for protein engineering to further extend lytic spectrum to Gram-negative bacteria and stabilize the protein.

## Materials & Methods

### Bacterial strains and human cell lines

All bacterial strains are presented in Table 3. *Neisseria* strains were plated from frozen aliquots onto chocolate agar plates (Difco GC Medium Base with BBL Hemoglobin Bovine (BD, USA) and Kellogg supplements (0.004% w/v glucose, 0.1 mg/mL glutamine, 0.2 μg/mL thiamine pyrophosphate, 0.5 μg/mL Fe(NO₃)₃·9H₂O)). *G. vaginalis* was cultured on Blood Lab Agar with 5% sheep blood (BioMaxima, Poland), *L. plantarum* on MRS plates, and *E. coli* strains on LB (BioMaxima, Poland). All strains were incubated with 5% CO_2_ (except for *E. coli*) at 37°C.

**Tab. 3.**
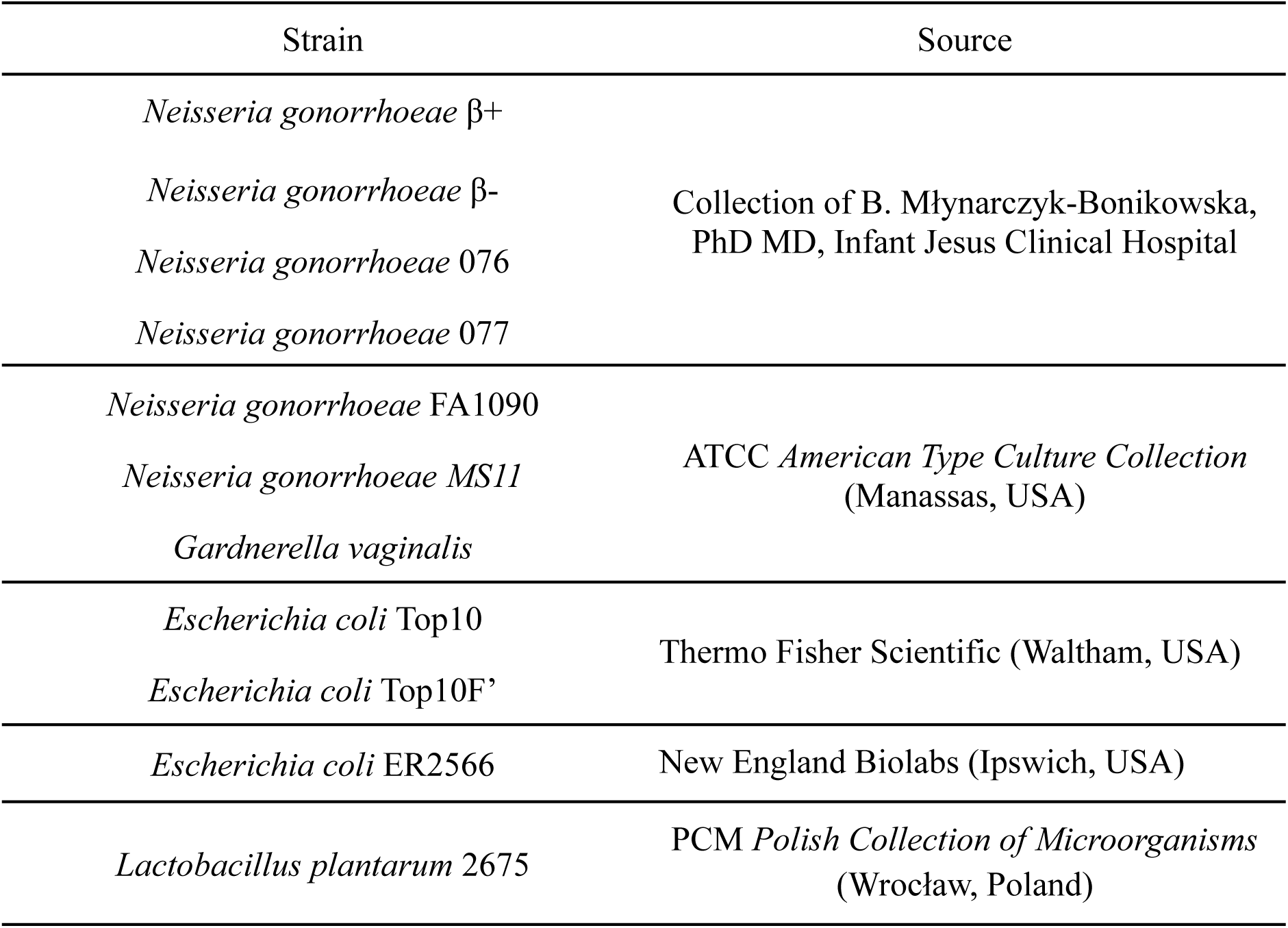
Bacterial strains used in this study.

ME-180 human cervical cell line (ATCC HTB-33) was cultured in McCoy medium (Biological Industries) supplemented with 10% heat-inactivated FBS (fetal bovine serum), antibiotics (100 U/mL penicillin + 100 μg/mL streptomycin) and 2.5 μg/mL amphotericin B (Biowest, France). Cells were incubated at 37°C with 5% CO_2_.

### Molecular cloning of potential prophage genes encoding endolysins

Primers for genes *ngo_0518* and *ngo_1649* introduced NheI and XhoI restriction sites (Table 4.). Start codons were omitted in primer design in order to introduce the 6x His-tag on the N-end of proteins of interest. PCR reaction was carried out using Phusion polymerase (Thermo Fisher Scientific, USA), with an annealing temperature of 71°C for both pairs of primers. Reaction parameters were set according to the producer’s manual, with one modification – twice the amount of reverse primer was added, as it has two annealing sites in the *N. gonorrhoeae* FA1090 template genome.

**Tab. 4.**
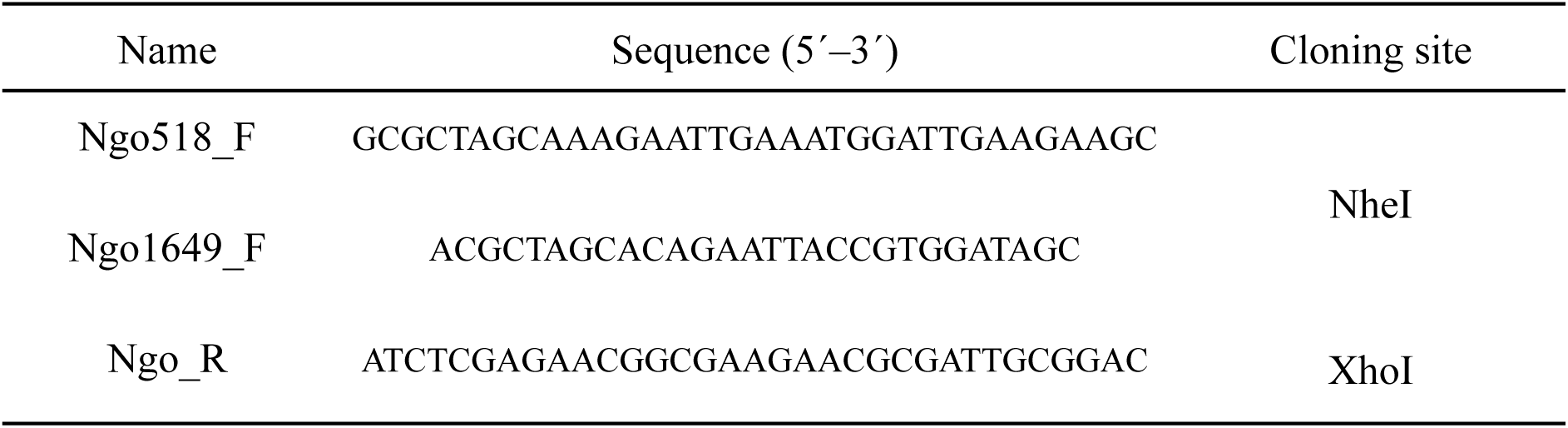
PCR primer sequences used in this study.

PCR products and pET-28a(+) vector plasmid (Novogene, China) were cut with NheI and XhoI enzymes (Thermo Fisher Scientific, USA) and ligated. After chemical transformation to *E. coli* Top 10 cells, plasmids were isolated and sequenced for proper cloning confirmation. We obtained two constructs: pET28a-518-His and pET28a-1649-His, carrying genes *ngo_0518* and *ngo_1649*, respectively.

### Protein expression and purification

For protein synthesis, *E. coli* ER2566 expression strain (NEB, USA) was transformed with pET28a-518-His or pET28a-1649-His plasmids and plated onto LB medium (BioMaxima, Poland) with kanamycin (final 35 μg/mL) and an additional 1% glucose. Overnight cultures of transformants were cultured in 100 mL of LB medium supplemented with kanamycin. They were induced at OD_600_ = 0.8 with 1 mM IPTG (3 h, 37°C, 180 rpm agitation). Cultures were centrifuged at 12 000 g for 20 min at 4°C. Pellet was resuspended in S buffer (PBS pH 7.4, 10 mM imidazole, 10% glycerol) supplemented with 1 mM PMSF. Cells were lysed by sonication (3 ✕ 5 min, amplitude 25%, pulse 30/30) and harvested by centrifugation under the same conditions as described above. 160 U of Viscolase (A&A Biotechnology, Poland) was added per 1 mL of supernatant, in order to reduce viscosity from nucleic acids release and increase purification quality. Protein synthesis efficacy was evaluated by SDS-PAGE gel electrophoresis.

Supernatants containing proteins were purified by affinity chromatography, using HisPur™ spin columns with Ni-NTA agarose resin (Thermo Fisher Scientific, USA). Columns were equilibrated with S buffer. Washing steps were performed using W1 buffer (S buffer with 25 mM imidazole) two times and W2 buffer (S buffer with 50 mM imidazole) once. Elution steps were performed with buffers E1 and E2 (S buffer containing 250 and 500 mM imidazole, respectively), resulting in a 5-step elution process. Each fraction was collected separately.

Fraction samples were analyzed by SDS-PAGE (Fig. S4.). Best fractions (selected based on purity of protein bands on the gel) were then further processed with dialysis in the D buffer (pH 8) containing 20 mM Tris-Cl, 250 mM NaCl and 10% glycerol. Dialysis was performed in nitrocellulose membrane sacs (Spectrum Laboratories Inc., Canada) for 24 h at 4°C. Purified protein samples were sterilized with 0.22 μm syringe filters and stored at -70°C. Concentration of purified proteins was measured using the Bradford technique with albumin standard (Thermo Fisher Scientific, USA). Samples were prepared with Bradford Reagent (Merck, Germany), according to the manual, and measured on a Sunrise spectrophotometer (Tecan, Switzerland). Protein concentration was 1.96 mg/mL and 1.6 mg/mL for Phi1gp518 and Phi3gp1649 fractions, respectively.

### Zymogram activity test

Peptidoglycan (PG) as substrate for zymogram was prepared from 0.5 g lyophilized *Micrococcus sp.* cells (Merck, Germany), suspended in 10 mL of MilliQ water and sterilized at ¾ atm. Zymogram SDS-PAGE stacking gels contained 4% polyacrylamide, and running gels were prepared with 15% polyacrylamide and 10% PG. All sample concentrations were set to 1 mg/mL. Lysozyme and mutanolysin (A&A Biotechnology, Poland) were used as positive controls. Protein and control samples were prepared with a 2X non-reducing sample buffer (100 mM Tris-Cl, pH 6.8, 20% glycerol and 0.1% bromophenol blue). Samples were incubated for 30 min on ice, then denatured for 3 min at 95°C. Two zymogram gels with identical sets of samples were run in a cooling room at 85V for 25 min, then 120V for 1 h. Gels were then washed four times with distilled water. To prevent any false positive results, one of the gels was immediately stained for 1.5 h at RT with staining solution (0.1% methylene blue, 0.01% KOH). Destaining at RT with distilled water (with gentle rocking) took about 1 h. The second gel was transferred for 30 min at RT to the renaturing buffer (25 mM Tris-Cl, pH 7.5, 0.1% Triton X-100, 150 mM NaCl, 10 mM MgCl_2_). After transfer into a fresh renaturing buffer, the gel was incubated at 37°C for 72 h with slow shaking (50 rpm) and then stained as described above. During destaining, clear bands would appear in spots where renatured proteins showed muralytic activity.

### Turbidity reduction assay

TRA (turbidity reduction assay) was performed using the protocol from Lavigne et al., 2004, adapted to *Neisseria* spp. Bacteria were cultured onto chocolate agar plates and incubated at 37°C with 5% CO_2_ for 24 h. Bacteria were then scraped, resuspended in sterile PBS, and OD_600_ was set to 0.6. Cells were pelleted by centrifugation at 1800 g, 4°C for 15 min. The total pellet was resuspended in a permeabilization buffer (7.4 pH PBS saturated with chloroform, ratio 1:1) – 40 mL per pellet from 100 mL of bacterial culture. Mixtures were incubated with gentle shaking (90 rpm) at 20°C for 25 min. Permeabilized cells were then centrifuged and washed with PBS, then centrifuged again. The pellet was then resuspended in PBS until OD_600_ reached 1.

30 μL of protein or control samples and 270 μL of permeabilized cells per well were added in triplicate to a 96-well plate. Final concentrations of endolysins were varied between experiments from 1 to 3000 nM. The dialysis buffer was used as a negative control. The plate was prepared on a cooling block and then placed immediately in a Sunrise spectrophotometer (Tecan, Switzerland). The plate was shaken for 10 s at low intensity, and kinetic measurement at 600 nm wavelength was performed with a 1-minute interval for 45 min at RT. Measurement data were analyzed with an activity calculator provided by Briers et al., 2007. For raw bacteria assays, we followed the same procedure, omitting the permeabilization step.

### Influence of pH and temperature on Phi1gp518 activity

This assay was based on the TRA test, described above. To assess pH range, permeabilized *N. gonorrhoeae* FA1090 cells were resuspended in eight different buffers (before adding endolysin): 20 mM citrate buffer (pH 3 – 5), PBS (pH 6 – 7), 20 mM Tris-Cl (pH 8 – 10). For temperature range evaluation, Phi1gp518 (final concentration 3000 nM) was incubated for 45 min at different temperatures: RT (on the bench), 30°C, 37°C, 40°C, 42°C, 50°C, 55°C, 60°C or 65°C.

### Biofilm assays

The methodology was previously adapted to gonococci (47). For all biofilm assays *N. gonorrhoeae* FA1090 was plated on chocolate agar plates and incubated at 37°C for 24 h with 5% CO_2_. Bacteria were then scraped and resuspended in liquid GC medium supplemented with Kellogg and 10 mM NaHCO_3_ until OD_600_ = 0.2. Biofilms were set up with non-permeabilized gonococci.

#### 1) Visualisation of gonococcal microcolony formation

A 96-well plate was set up as described above. Just before measurements, we added 3000 nM Phi1gp518 to six wells with bacteria, while six additional control wells remained untreated. The plate was installed on the JuLi Stage ™ imaging system (NanoEntek, Korea) in an incubator set to 37°C with 5% CO_2_. During a 12-hour-long incubation, bright light microscopic images with 40X magnification were taken in 1-hour intervals.

#### 2) Crystal violet assay on formed biofilm

Bacterial suspension (prepared as described above) was added in the volume of 200 μL per well in a 96-well plate (NEST Biotechnology). After 20 h incubation at 37°C with 5% CO_2_, supernatants were discarded, and each well was washed three times with 7.2 pH PBS. Protein samples (final 1000, 3000, or 6000 nM Phi1gp518) or control sample (PBS) were added in a volume of 50 μL per well to six technical replicates. Plates were incubated for 2 h at 37°C. Wells were washed with tap water and stained with 0.1% crystal violet for 15 min at RT. Plates were washed three times with tap water and left to dry overnight. 30% acetic acid was used to solubilize crystal violet, and the OD_550_ measurement was performed in a fresh 96-well plate.

#### 3) Effect of Phi1gp518 on formed biofilm

Biofilms were set up by adding 3 mL of bacterial suspension onto a plate with a glass bottom (Corning). After 20 h incubation at 37°C with 5% CO_2_, supernatants were discarded and each well was washed twice with PBS. Purified protein Phi1gp518 (3000 nM) or PBS as a control sample were added in a volume of 2 mL per plate. Plates were incubated for 2 h at 37°C, 5% CO_2_. After discarding supernatants, plates were washed twice with PBS. Biofilms were then stained with 1 mL of LIVE/DEAD ™ BacLight ™ (Invitrogen, USA) 1X solution and incubated for 20 min at RT in the dark. Plates were visualized by the Laboratory of Confocal Microscopy (University of Warsaw). Image analysis was performed using ImageJ (48).

### Evaluation of endolysin cytotoxicity on human cells

Two 96-well plates (NEST, China) were set up with 1 x 10^4^ ME-180 cells in 100 μL per well. After reaching confluence (4 x 10^4^ cells/well), the medium was discarded, and the cells were washed three times with PBS. Dilutions of protein and control samples were prepared in cell medium (with 5% FBS). Phi1gp518 (3000 nM, 6000 nM), 0.1% saponin, PBS, and dialysis buffer (10-fold dilution in cell medium) were added in six technical replicates (100 μL per well). Plates were incubated for 24 and 48 h at 37°C with 5% CO_2_. Cytotoxicity assay was performed using Cell Proliferation Kit I MTT (Roche), according to the manufacturer’s instructions. Numerical data were obtained with a Sunrise spectrophotometer (Tecan) at a 550 nm wavelength.

### Statistical analysis

All data were obtained from experiments designed with at least three technical replicates and at least two biological replicates. Numerical data were processed in the GraphPad Prism (version 8.0.2 for Windows, GraphPad Software, www.graphpad.com). Normality was assessed with the Shapiro-Wilk test. We used Brown-Forsythe and Welch ANOVA as a parametric test or Kruskal-Wallis as a non-parametric one. Statistical significance was determined as ‘ns’ (not significant), * (p < 0.05), ** (p < 0.01), *** (p < 0.001), or **** (p < 0.0001). This notation is consistent across all figures.

## Conflicts of interest

The authors declare no conflict of interest.

## Funding

This work was funded by the National Science Centre, Poland, grant no. 2021/43/O/NZ6/00379, received by Monika Adamczyk-Popławska, PhD.

## Acknowledgements

We would like to thank B. Młynarczyk-Bonikowska, PhD, MD for providing us with clinical gonococcal strains.

## Supplementary

**Fig. S1.**
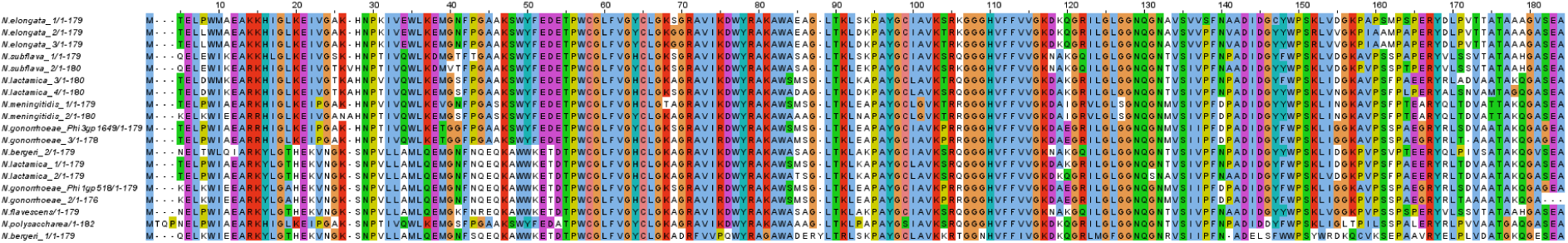
MAFFT multiple alignment of amino acid sequences of NlpC/P60 peptidases among *Neisseria* genus

**Fig. S2.**
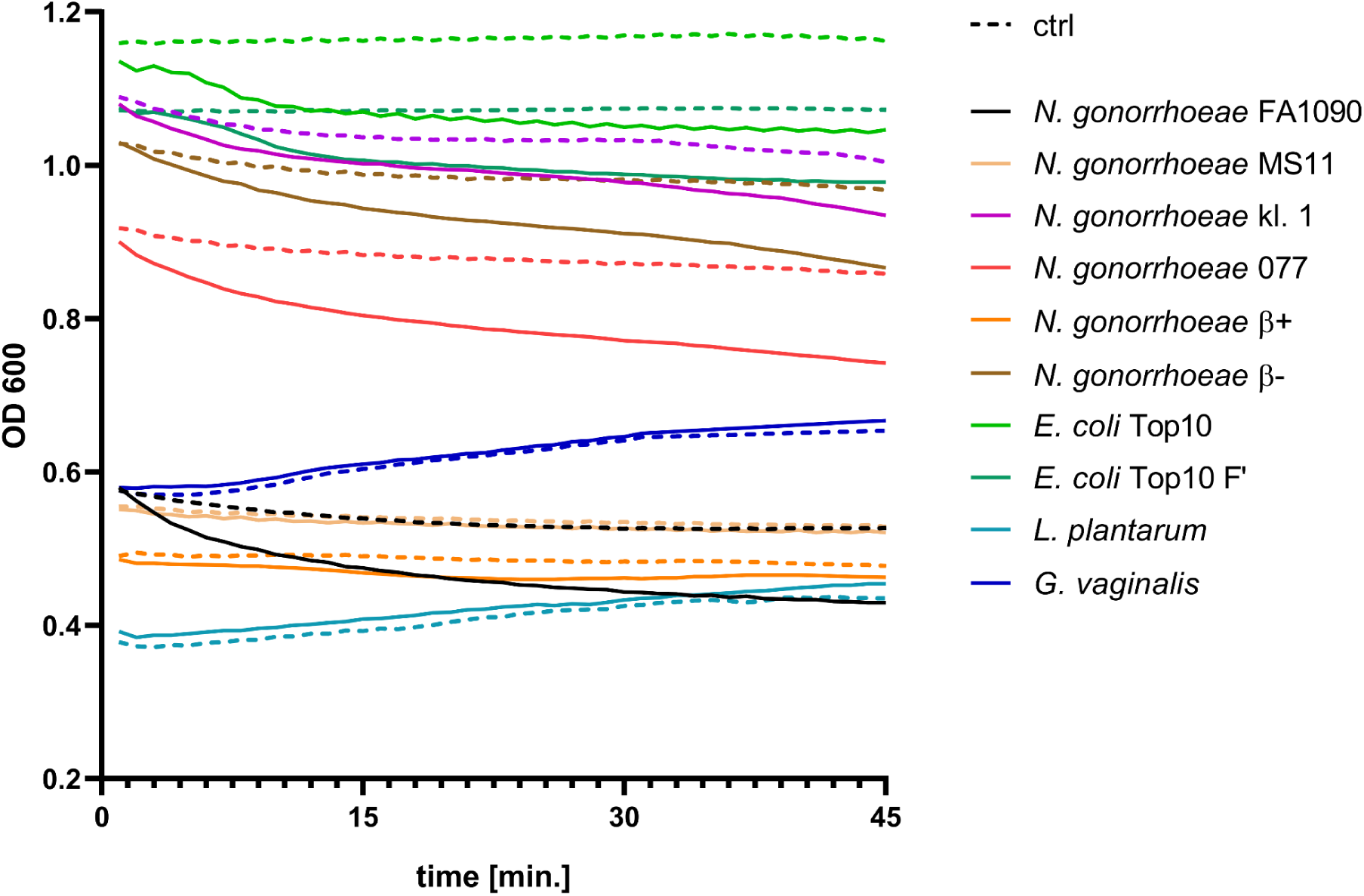
TRA with raw bacterial strains; ctrl: bacteria only, solid lines: bacteria treated with Phi1gp518

**Fig. S3.**
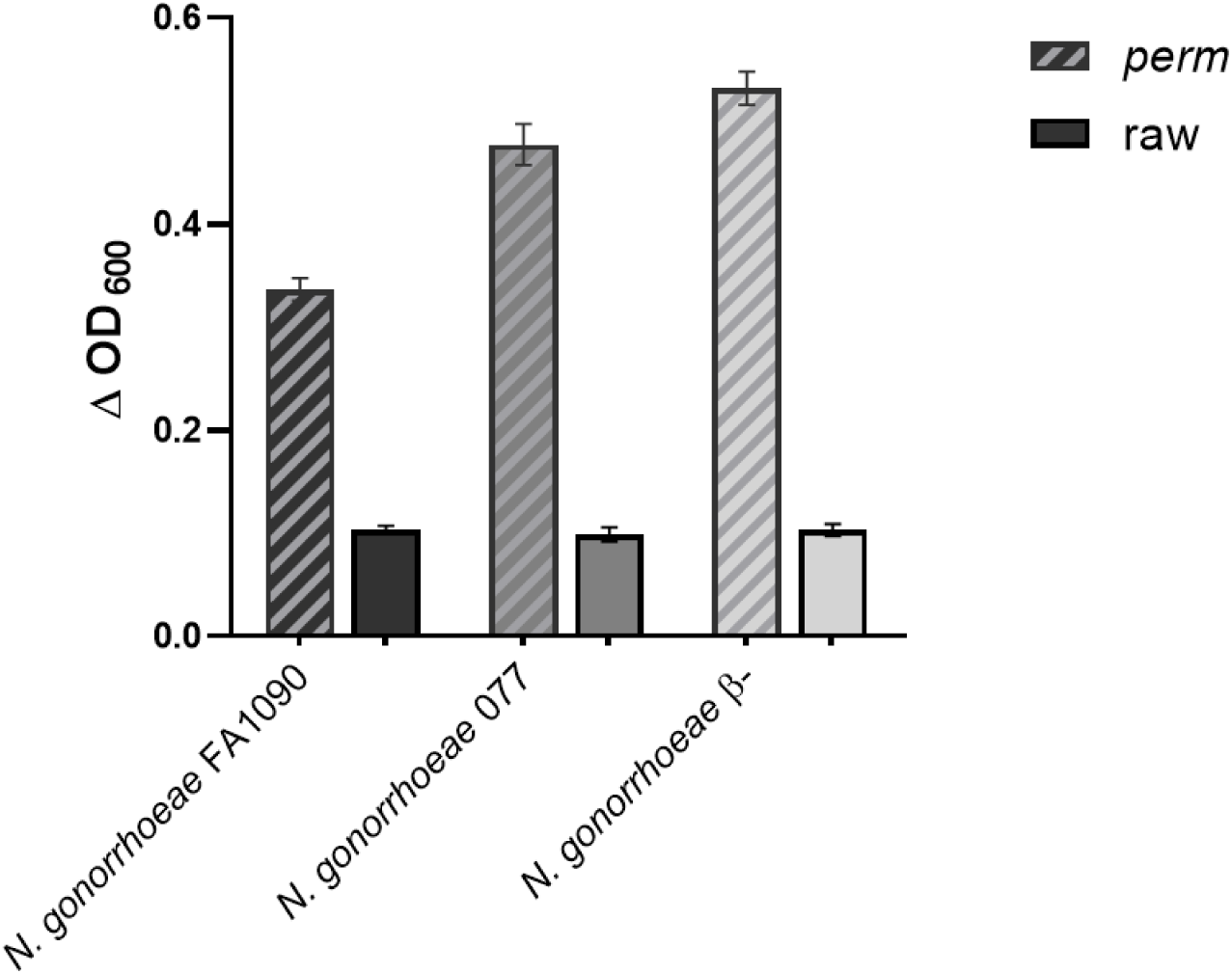
Comparison between absolute OD_600_ decrease of permeabilized gonococcal strains; Δ OD_600_ value was calculated as difference between OD_600_ at starting and ending time point of measurement, with subtraction of control (untreated bacteria) value for each strain

**Fig. S4.**
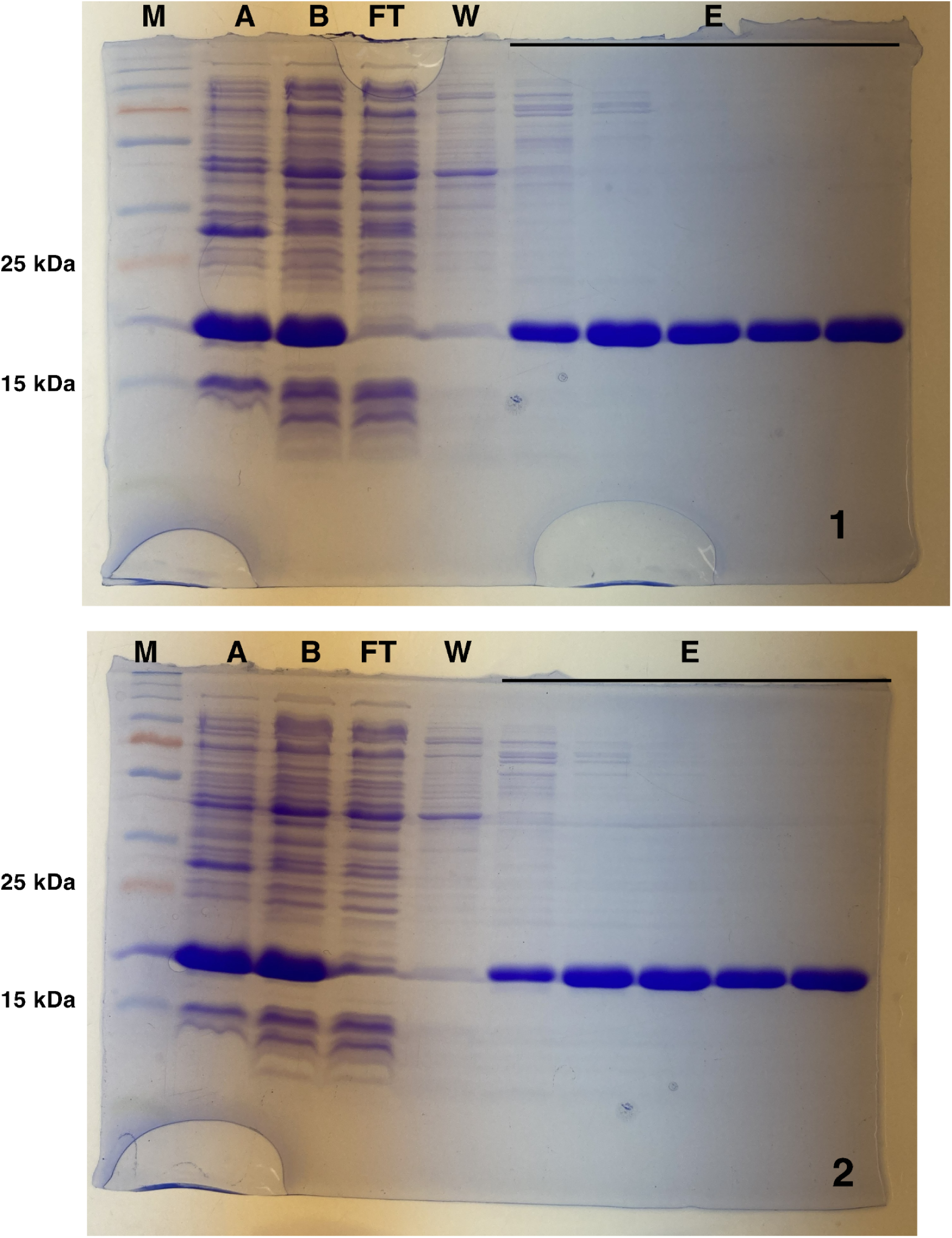
SDS-PAGE gel images showing different fractions collected during purification of (1) Phi1gp518 or (2) Phi3gp1649 proteins; lanes include samples of (M) protein size marker, (A) induction pellet, (B) crude extract, (FT) flowthrough, (W) first wash, (E) elution rounds

## References

1. Abara WE, Kirkcaldy RD, Bernstein KT, Galloway E, Learner ER. 2025. Effectiveness of MenB-4C Vaccine Against Gonorrhea: A Systematic Review and Meta-analysis. J Infect Dis 231:61–70.

2. Quillin SJ, Seifert HS. 2018. Neisseria gonorrhoeae host adaptation and pathogenesis. Nat Rev Microbiol 16:226–240.

3. http://who.int/news-room/fact-sheets.

4. Adamczyk-Popławska M, Golec P, Piekarowicz A, Kwiatek A. 2024. The potential for bacteriophages and prophage elements in fighting and preventing the gonorrhea. Crit Rev Microbiol 50:769–784.

5. Roach DR, Donovan DM. 2015. Antimicrobial bacteriophage-derived proteins and therapeutic applications. Bacteriophage 5:e1062590.

6. Latka A, Maciejewska B, Majkowska-Skrobek G, Briers Y, Drulis-Kawa Z. 2017. Bacteriophage-encoded virion-associated enzymes to overcome the carbohydrate barriers during the infection process. Appl Microbiol Biotechnol 101:3103–3119.

7. Gilmer DB, Schmitz JE, Euler CW, Fischetti VA. 2013. Novel bacteriophage lysin with broad lytic activity protects against mixed infection by Streptococcus pyogenes and methicillin-resistant Staphylococcus aureus. Antimicrob Agents Chemother 57:2743–2750.

8. Oh J, Warner M, Ambler JE, Schuch R. 2023. The Lysin Exebacase Has a Low Propensity for Resistance Development in Staphylococcus aureus and Suppresses the Emergence of Resistance to Antistaphylococcal Antibiotics. Microbiol Spectr 11:e0526122.

9. Laumen JGE, Abdellati S, Manoharan-Basil SS, Van Dijck C, Van den Bossche D, De Baetselier I, de Block T, Malhotra-Kumar S, Soentjes P, Pirnay J-P, Kenyon C, Merabishvili M. 2022. Screening of Anorectal and Oropharyngeal Samples Fails to Detect Bacteriophages Infecting Neisseria gonorrhoeae. Antibiotics 11:268.

10. Orazi G, Collins AJ, Whitaker RJ. 2022. Prediction of Prophages and Their Host Ranges in Pathogenic and Commensal Neisseria Species. mSystems 7:e0008322.

11. Shaskolskiy B, Kravtsov D, Kandinov I, Dementieva E, Gryadunov D. 2022. Genomic Diversity and Chromosomal Rearrangements in Neisseria gonorrhoeae and Neisseria meningitidis. Int J Mol Sci 23:15644.

12. Piekarowicz A, Majchrzak M, Kłyz A, Adamczyk-Popławska M. 2006. Analysis of the filamentous bacteriophage genomes integrated into Neisseria gonorrhoeae FA1090 chromosome. Pol J Microbiol 55:251–260.

13. Piekarowicz A, Kłyz A, Majchrzak M, Adamczyk-Popławska M, Maugel TK, Stein DC. 2007. Characterization of the dsDNA prophage sequences in the genome of Neisseria gonorrhoeae and visualization of productive bacteriophage. BMC Microbiol 7:66.

14. Griffin ME, Klupt S, Espinosa J, Hang HC. 2023. Peptidoglycan NlpC/P60 peptidases in bacterial physiology and host interactions. Cell Chem Biol 30:436–456.

15. Wishart DS, Han S, Saha S, Oler E, Peters H, Grant JR, Stothard P, Gautam V. 2023. PHASTEST: faster than PHASTER, better than PHAST. Nucleic Acids Res 51:W443–W450.

16. Olson RD, Assaf R, Brettin T, Conrad N, Cucinell C, Davis JJ, Dempsey DM, Dickerman A, Dietrich EM, Kenyon RW, Kuscuoglu M, Lefkowitz EJ, Lu J, Machi D, Macken C, Mao C, Niewiadomska A, Nguyen M, Olsen GJ, Overbeek JC, Parrello B, Parrello V, Porter JS, Pusch GD, Shukla M, Singh I, Stewart L, Tan G, Thomas C, VanOeffelen M, Vonstein V, Wallace ZS, Warren AS, Wattam AR, Xia F, Yoo H, Zhang Y, Zmasek CM, Scheuermann RH, Stevens RL. 2023. Introducing the Bacterial and Viral Bioinformatics Resource Center (BV-BRC): a resource combining PATRIC, IRD and ViPR. Nucleic Acids Res 51:D678–D689.

17. Altschul SF, Gish W, Miller W, Myers EW, Lipman DJ. 1990. Basic local alignment search tool. J Mol Biol 215:403–410.

18. Zimmermann L, Stephens A, Nam S-Z, Rau D, Kübler J, Lozajic M, Gabler F, Söding J, Lupas AN, Alva V. 2018. A Completely Reimplemented MPI Bioinformatics Toolkit with a New HHpred Server at its Core. J Mol Biol 430:2237–2243.

19. Gabler F, Nam S-Z, Till S, Mirdita M, Steinegger M, Söding J, Lupas AN, Alva V. 2020. Protein Sequence Analysis Using the MPI Bioinformatics Toolkit. Curr Protoc Bioinforma 72:e108.

20. Pérez Medina KM, Dillard JP. 2018. Antibiotic Targets in Gonococcal Cell Wall Metabolism. Antibiotics 7:64.

21. https://zhanggroup.org//TM-align/.

22. Jumper J, Evans R, Pritzel A, Green T, Figurnov M, Ronneberger O, Tunyasuvunakool K, Bates R, Žídek A, Potapenko A, Bridgland A, Meyer C, Kohl SAA, Ballard AJ, Cowie A, Romera-Paredes B, Nikolov S, Jain R, Adler J, Back T, Petersen S, Reiman D, Clancy E, Zielinski M, Steinegger M, Pacholska M, Berghammer T, Bodenstein S, Silver D, Vinyals O, Senior AW, Kavukcuoglu K, Kohli P, Hassabis D. 2021. Highly accurate protein structure prediction with AlphaFold. Nature 596:583–589.

23. Anantharaman V, Aravind L. 2003. Evolutionary history, structural features and biochemical diversity of the NlpC/P60 superfamily of enzymes. Genome Biol 4:R11.

24. Katoh K, Misawa K, Kuma K, Miyata T. 2002. MAFFT: a novel method for rapid multiple sequence alignment based on fast Fourier transform. Nucleic Acids Res 30:3059–3066.

25. Tisza MJ, Buck CB. 2021. A catalog of tens of thousands of viruses from human metagenomes reveals hidden associations with chronic diseases. Proc Natl Acad Sci U S A 118:e2023202118.

26. Escobar CA, Cross TA. 2018. False positives in using the zymogram assay for identification of peptidoglycan hydrolases. Anal Biochem 543:162–166.

27. Lavigne R, Briers Y, Hertveldt K, Robben J, Volckaert G. 2004. Identification and characterization of a highly thermostable bacteriophage lysozyme. Cell Mol Life Sci 61:2753–2759.

28. Briers Y, Lavigne R, Volckaert G, Hertveldt K. 2007. A standardized approach for accurate quantification of murein hydrolase activity in high-throughput assays. J Biochem Biophys Methods 70:531–533.

29. Szadkowska M, Olewniczak M, Kloska A, Jankowska E, Kapusta M, Rybak B, Wyrzykowski D, Zmudzinska W, Gieldon A, Kocot A, Kaczorowska A-K, Nierzwicki L, Makowska J, Kaczorowski T, Plotka M. 2022. A Novel Cryptic Clostridial Peptide That Kills Bacteria by a Cell Membrane Permeabilization Mechanism. Microbiol Spectr 10:e0165722.

30. Chen X, Lu Y, Chen T, Li R. 2021. The Female Vaginal Microbiome in Health and Bacterial Vaginosis. Front Cell Infect Microbiol 11:631972.

31. Pinto D, Gonçalo R, Louro M, Silva MS, Hernandez G, Cordeiro TN, Cordeiro C, São-José C. 2022. On the Occurrence and Multimerization of Two-Polypeptide Phage Endolysins Encoded in Single Genes. Microbiol Spectr 10:e0103722.

32. McKerral JC, Papudeshi B, Inglis LK, Roach MJ, Decewicz P, McNair K, Luque A, Dinsdale EA, Edwards RA. 2023. The Promise and Pitfalls of Prophages. 2023.04.20.537752. bioRxiv: The Preprint Server for Biology 10.1101/2023.04.20.537752.

33. Gangakhedkar R, Jain V. 2024. Elucidating the molecular properties and anti-mycobacterial activity of cysteine peptidase domain of D29 mycobacteriophage endolysin. J Virol 98:e01328–24.

34. Rosenthal RS, Wright RM, Sinha RK. 1980. Extent of peptide cross-linking in the peptidoglycan of Neisseria gonorrhoeae. Infect Immun 28:867–875.

35. Gutiérrez D, Briers Y. 2021. Lysins breaking down the walls of Gram-negative bacteria, no longer a no-go. Curr Opin Biotechnol 68:15–22.

36. O’Hanlon DE, Moench TR, Cone RA. 2013. Vaginal pH and microbicidal lactic acid when lactobacilli dominate the microbiota. PloS One 8:e80074.

37. Leal J, Smyth HDC, Ghosh D. 2017. Physicochemical properties of mucus and their impact on transmucosal drug delivery. Int J Pharm 532:555–572.

38. Çakırgöz Ç, Çakırgöz E, Kirmiç D, Hasdemir PS, Değerli K. 2025. The impact of serum estradiol levels on vaginal pH and Candida infections during infertility treatment. Steroids 221:109655.

39. Manca B, Buffi G, Magri G, Del Vecchio M, Taddei AR, Pezzicoli A, Giuliani M. 2023. Functional characterization of the gonococcal polyphosphate pseudo-capsule. PLOS Pathog 19:e1011400.

40. Antonova NP, Vasina DV, Lendel AM, Usachev EV, Makarov VV, Gintsburg AL, Tkachuk AP, Gushchin VA. 2019. Broad Bactericidal Activity of the Myoviridae Bacteriophage Lysins LysAm24, LysECD7, and LysSi3 against Gram-Negative ESKAPE Pathogens. Viruses 11:284.

41. Vollmer W, Bertsche U. 2008. Murein (peptidoglycan) structure, architecture and biosynthesis in Escherichia coli. Biochim Biophys Acta 1778:1714–1734.

42. Gökmen GG, Sarıyıldız S, Cholakov R, Nalbantsoy A, Baler B, Aslan E, Düzel A, Sargın S, Göksungur Y, Kışla D. 2024. A novel Lactiplantibacillus plantarum strain: probiotic properties and optimization of the growth conditions by response surface methodology. World J Microbiol Biotechnol 40:66.

43. Janulaitiene M, Gegzna V, Baranauskiene L, Bulavaitė A, Simanavicius M, Pleckaityte M. 2018. Phenotypic characterization of Gardnerella vaginalis subgroups suggests differences in their virulence potential. PloS One 13:e0200625.

44. Petrova OE, Sauer K. 2016. Escaping the biofilm in more than one way: desorption, detachment or dispersion. Curr Opin Microbiol 30:67–78.

45. Schaub RE, Dillard JP. 2019. The Pathogenic Neisseria Use a Streamlined Set of Peptidoglycan Degradation Proteins for Peptidoglycan Remodeling, Recycling, and Toxic Fragment Release. Front Microbiol 10:73.

46. Gerstmans H, Criel B, Briers Y. 2018. Synthetic biology of modular endolysins. Biotechnol Adv 36:624–640.

47. Kwiatek A, Mrozek A, Bacal P, Piekarowicz A, Adamczyk-Popławska M. 2015. Type III Methyltransferase M.NgoAX from Neisseria gonorrhoeae FA1090 Regulates Biofilm Formation and Interactions with Human Cells. Front Microbiol 6:1426.

48. Schneider CA, Rasband WS, Eliceiri KW. 2012. NIH Image to ImageJ: 25 years of image analysis. Nat Methods 9:671–675.

